# A GATA6-centered gene regulatory network involving HNFs and ΔNp63 controls plasticity and immune escape in pancreatic cancer

**DOI:** 10.1101/2020.04.09.033456

**Authors:** Bernhard Kloesch, Natascha Hruschka, Vivien Ionasz, Rupert Oellinger, Sebastian Mueller, Jaime Martinez de Villareal, Hans-Peter Dienes, Martin Schindl, Elisabeth Gruber, Judith Stift, Gwen Lomberk, Barbara Seidler, Dieter Saur, Roland Rad, Raul Urrutia, Francisco X. Real, Paola Martinelli

**Author notes:** Corresponding author,. Current address: Boehringer-Ingelheim RCV, Vienna, Austria.

## Abstract

**Objective:** Molecular taxonomy of tumors is the foundation of personalized medicine and is becoming of paramount importance for therapeutic purposes. Four transcriptomics-based classification systems of pancreatic ductal adenocarcinoma (PDAC) exist, which consistently identified a subtype of highly aggressive PDACs with basal-like features, including ΔNp63 expression and loss of the epithelial master regulator GATA6. We investigated the precise molecular events driving PDAC progression and the emergence of the basal program.

**Design:** We combined the analysis of patient-derived transcriptomics datasets and tissue samples with mechanistic experiments using a novel dual-recombinase mouse model for Gata6 deletion at late stages of KRas^G12D^-driven pancreatic tumorigenesis (Gata6^LateKO^).

**Results:** This comprehensive human-to-mouse approach allowed us to show that GATA6 loss is necessary, but not sufficient, for the expression of a basal program in patients and in mice. The concomitant loss of HNF1A and HNF4A, likely through epigenetic silencing, is required for the full phenotype switch. Moreover, Gata6 deletion in mice dramatically increased the metastatic rate, with a propensity for lung metastases. Through RNA-Seq analysis of primary cells isolated from mouse tumors, we show that Gata6 inhibits tumor cell plasticity and immune evasion, suggesting that it works as a barrier for acquiring the fully developed basal and metastatic phenotype.

**Conclusions:** Our work provides both a mechanistic molecular link between the basal phenotype and metastasis and a valuable preclinical tool to investigate the most aggressive subtype of PDAC. These data, therefore, are important for understanding the pathobiological features underlying the heterogeneity of pancreatic cancer in both mice and human.

**What is already known about this subject?:**

- Multiple transcriptomics-based studies have identified a basal-like subtype of pancreatic ductal adenocarcinoma (PDAC) with especially poor prognosis.
- Loss of GATA6 in PDAC cells is associated with altered differentiation, including ectopic expression of basal markers such as KRT14.
- Aberrant expression of the ΔNp63 transcription factor can drive the expression of the basal transcriptional program.

**What are the new findings?:**

- Loss of GATA6 expression is necessary but not sufficient for the expression of ΔNp63 and the basal phenotype.
- Concomitant silencing of HNF4A and HNF1A, possibly through epigenetic mechanisms, is required for the full-blown phenotype.
- *Gata6* deletion in established murine tumors favors the basal and metastatic phenotype, with a lung tropism, in a next-generation model of KRas^G12D^-driven PDAC.
- Loss of GATA6 expression is associated with features of immune escape in mouse and human PDAC cells.

**How might it impact on clinical practice in the foreseeable future?:**

- The combined analysis of GATA6, HNFs, and TP63 expression in patient-derived samples will provide a more precise classification of PDAC.
- Restoration of the classical PDAC phenotype may not only reduce metastatic potential but also increase immune recognition of tumor cells.

## INTRODUCTION

Molecular taxonomy of tumors harbors great potential for the development of personalized medicine. In the case of pancreatic ductal adenocarcinoma (PDAC), molecular classification is still in the early days but has already revealed the existence of multiple subtypes with distinctive biological and clinical behavior and likely specific vulnerabilities. Four PDAC taxonomies were proposed until now, differing in the number of subgroups and nomenclature^1–4^. Despite these discrepancies, all classifications identified a PDAC subtype with loss of cell identity features, associated with significantly worse survival in patients. This subtype, called “quasi-mesenchymal”^2^, “basal-like”^3^, “squamous”^1^, “pure basal”^4^, or “basal A/B”^5^ showed rather homogeneous gene expression profiles across classifications^6^. We will refer to this PDAC subtype as “basal”. A better understanding of the molecular drivers of this aggressive PDAC subtype would improve patients’ management in the context of an almost invariably lethal malignancy.

The transcription factor GATA6, a crucial regulator of acinar cell differentiation^7^ and suppressor of KRas^G12V^-driven tumorigenesis in mice^8^, was highly expressed in the classical subtype^2^ and silenced through promoter methylation in squamous tumors^1^. We confirmed that GATA6 was lost in a subset of PDACs, in association with a basal-like differentiation, and shed light on the underlying mechanism^9^. GATA6 silencing resulted in an epithelial-to-epithelial transition (E^2^T) and an epithelial-to-mesenchymal transition (EMT) while its overexpression induced mesenchymal-to-epithelial transition (MET), supporting its central role in determining the phenotype of PDACs^8 9^. Consistently, loss of GATA6 - as a single biomarker - identified basal tumors as efficiently as the corresponding gene signature in a cohort of metastatic PDAC patients^10^.

Recent publications indicated that the basal phenotype is driven by broad epigenomic reprogramming, especially at superenhancers, controlled by ΔNp63^11–13^, the shorter isoform of the TP63 transcription factor marking the basal layer of stratified epithelia. Interestingly, GATA6 itself was identified as being controlled by a superenhancer lost in basal patient-derived cells^14^, suggesting that loss of GATA6 might be embedded in the basal program rather than driving it.

Here we aimed at elucidating whether loss of GATA6 is the cause or the consequence of the basal phenotype in PDAC. By combining the analysis of patient-derived samples and transcriptomics datasets, *in vitro* experiments with PDAC cells, and a next-generation *KRas^G12D-^*driven mouse PDAC model where *Gata6* was deleted at late stages of tumorigenesis, we show that GATA6 loss is necessary, but not sufficient, for the appearance of a basal program in PDAC. Concomitant downregulation of HNF1A and HNF4A is required for the full phenotypic switch. Additionally, *Gata6* loss in high-grade preneoplastic lesions (PanINs) favored the development of metastases in mice, possibly by promoting plasticity and immune escape of tumor cells. We demonstrate that an epithelial/progenitor transcriptional network acts as a barrier against tumor progression, and provide a molecular link between the basal gene program *in vivo* and the metastatic spread in PDAC.

## METHODS & MATERIAL

All relevant methods and materials can be found in the online supplement.

## RESULTS

### GATA6 loss is necessary for the expression of the basal program

We analyzed five PDAC transcriptomic datasets with molecular classification, which revealed that GATA6 expression was consistently lower in the poorly differentiated subtypes (quasi-mesenchymal^2^ P=0.008, basal-like^3^ P=5.54e-10, squamous^1^ P=1.57e-11, pure-basal^4^ P<2e-16, Basal A/B^5^ P<0.0001) (Figure 1A). Since ΔNp63 was suggested to drive the basal transcriptional program in PDAC^11 12^, we explored its relationship with GATA6 in 4/5 of the datasets (the Collisson was excluded due to low sample size) plus the TCGA PAAD dataset. TP63 expression was negatively correlated with GATA6 expression in 4/5 datasets (Figure 1B, Supplementary Figure 1A). Additionally, ΔNp63-target genes^15^ were significantly enriched among those upregulated in GATA6^low^ tumors (bottom quartile) in 3/5 datasets and showed a tendency in the remaining 2 (Figure 1C, Supplementary Figure 1B), supporting that the ΔNp63-dependent program is induced when GATA6 is lost. We showed previously that GATA6 loss in PDAC associates with ectopic expression of the basal marker KRT14 in a small collection of patient-derived samples^9^. We measured GATA6, TP63, and KRT14 expression with IHC in an independent larger set of 60 formalin-fixed paraffin-embedded (FFPE) tissues from PDAC resections (Figure 1D). GATA6 expression was lost broadly or focally in 23/60 patients (38.3%). In addition, KRT14 was exclusively expressed in GATA6^low^ tumors (16/23, 69,6% P=1.64e^-09^) and TP63 expression was detected in 14/23 (60.1%) GATA6^low^ and 10/37 (27%) GATA6^high^ tumors (P=0.014) (Supplementary Figure 2A). Of note, the GATA6^high^/TP63^pos^ tumors only had small foci of TP63-positive cells, which, upon more detailed analysis, were found to be located in metaplastic lesions containing GATA6-negative cells in 9/10 cases (Supplementary Figure 2B, black arrowhead). Moreover, we compared the frequency of basal phenotypes between the top and bottom GATA6 expression quartiles (GATA6^high^, GATA6^low^) in the five PDAC datasets with classification and observed only 2/207 GATA6^high^/basal cases (Supplementary Figure 1C). These data strongly indicate that GATA6 loss is necessary for the expression of the basal phenotype.

**Figure 1.**
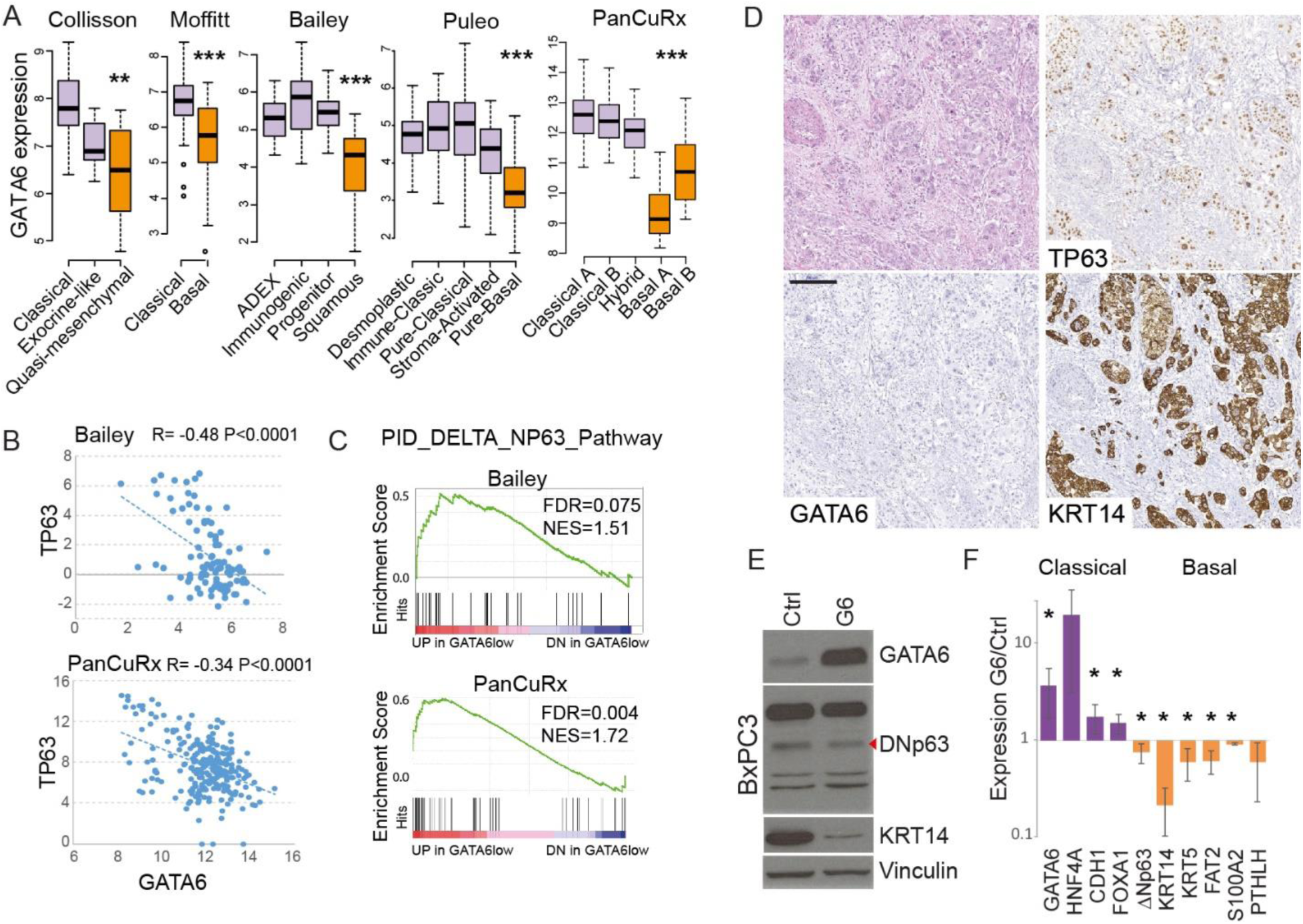
GATA6 loss is necessary for the expression of the basal-like program. A) Analysis of GATA6 mRNA expression in PDAC datasets with transcriptomics-based molecular classification. B) Correlation between TP63 and GATA6 mRNA expression in the indicated PDAC datasets. C) Enrichment of the gene set “ ΔNp63 target genes” among the genes up-regulated in GATA6^low^ versus GATA6^high^ tumors in the indicated datasets. D) Representative images of basal human PDAC. H&E (top left) and immunohistochemical stainings for TP63, GATA6, and KRT14. Scale bar = 200µm. E) Expression of GATA6, ΔNp63 (red arrowhead), and KRT14 in control (Ctrl) and GATA6-overexpressing (G6) BxPC3 cells, analyzed by western blotting of whole protein lysates. F) Expression of a set of classical and basal genes in GATA6-overexpressing BxPC3 cells compared to Ctrl cells, measured by RT-qPCR. Results are shown as mean ± standard deviation of at least n=3 biological replicates. *P>0.05.

To understand the hierarchical relationship between GATA6 and ΔNp63 in the regulation of the basal phenotype, we re-expressed GATA6 in BxPC3, a PDAC cell line with high levels of ΔNp63^11 12^. GATA6 overexpression led to a 40% reduction of ΔNp63 protein (P=4.15e-04) and 30% reduction of the mRNA (P=0.03) (Figure 1E, Supplementary Figure 2C). Consistently, using qRT-PCR we observed the up-regulation of classical (HNF4A, CDH1, FOXA1) and down-regulation of basal markers (ΔNp63, KRT14, KRT5, FAT2, S100A2, PTHLH); KRT14 was strongly reduced both at mRNA (80%, P=6.3e-06) and protein level (80%, P=1.42e-06) (Figure 1E, Supplementary Figure 2C) (Figure 1F). Intriguingly, we did not observe clear changes in proliferation, migration, or matrigel invasion *in vitro* (not shown).

A re-analysis of published RNA-Seq data^12^ showed that TP63 knock-out in BxPC3 cells significantly induced GATA6 expression (Adj.P=0.02) while ΔNp63 overexpression in PaTu8988S cells only resulted in a small, not significant, decrease in GATA6 mRNA (Supplementary Figure 2D). ChIP-Seq showed a TP63 peak downstream of GATA6 TSS (not shown), possibly indicating a direct repression. The intersection between TP63 ChIP-Seq peaks in BxPC3 from the same report^12^ and GATA6 ChIP-Seq peaks from our previous work showed limited overlap (0.8% of GATA6 peaks, 10.7% of TP63 peaks), suggesting that the two transcription factors control separate programs in basal vs classical cells. Interestingly, only 15 out of 41036 GATA6 peaks were located on regions identified as “Squamous elements” by Somerville et al^12^ (Supplementary Figure 2E). These data indicate that, while important, neither GATA6 loss nor ΔNp63 expression is sufficient to drive a full phenotypic switch in PDAC cells.

### GATA6 cooperates with HNF1A and HNF4A to sustain the classical phenotype

To identify the crucial molecular events downstream of GATA6 loss, we analyzed the PanCuRx dataset, including the largest series of all-stages PDAC samples and thus better representing the PDAC patient population than datasets only including resectable tumors. We compared GATA6^low^/Basal versus GATA6^low^/Classical tumors. GSEA revealed that HNF1A and HNF4A putative target genes were enriched among the upregulated transcripts in GATA6^low^/Classical tumors (Figure 2A). Accordingly, HNF1A and HNF4A mRNAs were significantly higher in GATA6^low^/Classical tumors, compared to the basal ones (Figure 2B and Supplementary Figure 3). Similarly, KRT14^pos^ regions of patients’ tumors showed reduced HNF4A protein levels, compared to KRT14^neg^ regions (Figure 2C).

**Figure 2.**
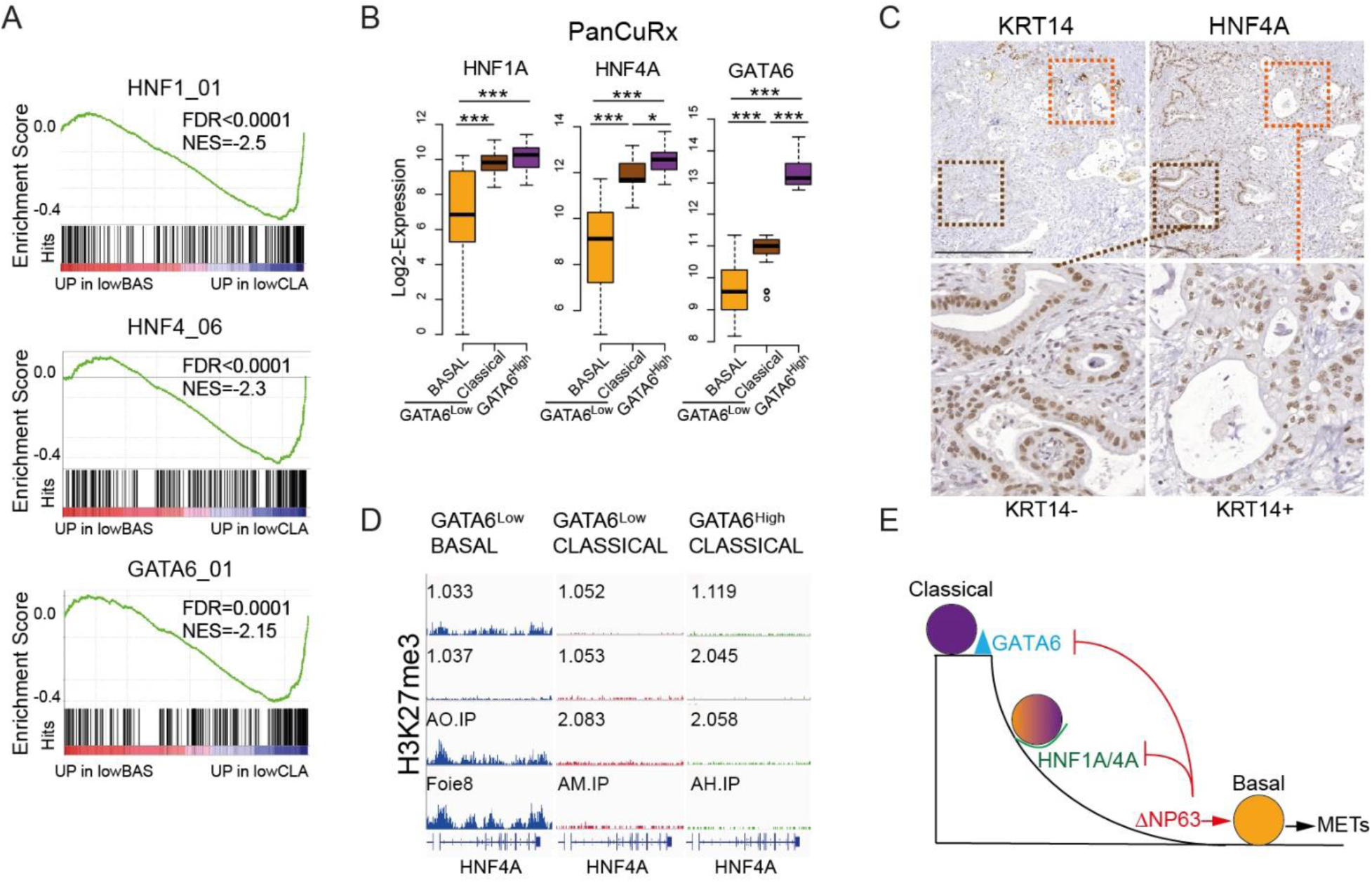
Concomitant loss of HNF1A and HNF4A is required for the expression of the basal phenotype after loss of GATA6. A) Enrichment plot of the gene sets containing putative HNF1A, HNF4A, and GATA6 target genes when comparing basal (lowBAS) and non-basal (lowCLA) GATA6^low^ tumors of the PanCuRx cohort. NES=Normalized Enrichment Score. B) Expression of HNF1A, HNF4A, and GATA6 in the different groups of patients in the PanCuRx dataset. *P<0.05, **P<0.001, ***P<0.0001. C) Representative images of KRT14 and HNF4A staining in a PDAC sample. Bottom images show KRT14^neg^/HNF4A^high^ (brown box) and KRT14^pos^/HNF4^low^ (orange box) regions. Scale bar: 500µM. D) H3K27me3 distribution along the *HNF4A* locus in PDX-derived cell lines of the three categories. E) The proposed model: GATA6 is the primary gatekeeper of the classical phenotype; HNF1A and HNF4A can block the full basal program but, once lost, the ΔNp63-driven basal program is fully expressed and drives PDAC progression toward metastasis. A negative feedback regulation driven by ΔNp63 might contribute to stabilize the basal phenotype.

We previously reported the most comprehensive epigenomics data available for a collection of basal and classical PDX-derived cell lines^14^. We reprocessed raw data to compare the epigenetic marks over *GATA6*, *HNF1A*, and *HNF4A* loci in GATA6^low^/Basal (n=4), GATA6^low^/Classical (n=5), and GATA6^high^/Classical (n=5) cell lines. Notably, while we found a marked accumulation of the heterochromatin marker H3K27me3 on the *HNF4A* locus in Gata6^low^/Basal cells, the locus was not epigenetically silenced in Gata6^low^/Classical ones (Figure 2D, Supplementary Figure 4A). *GATA6* showed a similar pattern but H3K27me3 was predominantly enriched upstream of the TSS (Supplementary Figure 4A). *HNF1A* was not highly marked with H3K27me3 in GATA6^low^/Basal cells (Supplementary Figure 4A).

H3K27ac and H3K4me3 patterns around the TSS of all the three genes were consistent with higher transcription in GATA6^low^/Classical and Gata6^high^/Classical cells, i.e. enrichment of these two markers of active chromatin was low or absent in GATA6^low^/Basal, with the exception of 1.037 cells, while it was high in all other cells (Supplementary Figure 4B-C). These data suggest that GATA6 and HNFs are epigenetically silenced in basal cells, while classical cells retain HNFs expression even when GATA6 is low. Importantly, although a subset of GATA6^high^/HNF1A^low^ and GATA6^high^/HNF4A^low^ tumors was present in all patient-derived datasets, none of those tumors was basal, further supporting that the coexisting loss of GATA6 and HNFs is required for the basal phenotype to emerge.

### Development of a next-generation mouse model to delete *Gata6* in established tumors

We showed previously that *Gata6* deletion at tumor initiation accelerates KRas^G12V^-driven pancreatic tumorigenesis^8^. However, *KRas^G12V^; Gata6^P-/-^* mice developed tumors that were generally well differentiated and Krt14-negative (not shown). To discriminate the effects related to tumor initiation from those related to tumor progression, we turned to a next-generation mouse model. For this purpose, we bred *Gata6^lox/lox^* mice^16^ with the dual recombinase mice harboring the *Pdx1-Flp*, *FSF-KRas^G12D^*, *FSF-R26^CreERT2^* ^17^ and *R26^Dual^* ^18^ alleles, to generate KFC mice (KRas, Flp, Cre). This new model allows the uncoupling between the activation of KRas^G12D^ expression and *Gata6* deletion (Figure 3A).

**Figure 3.**
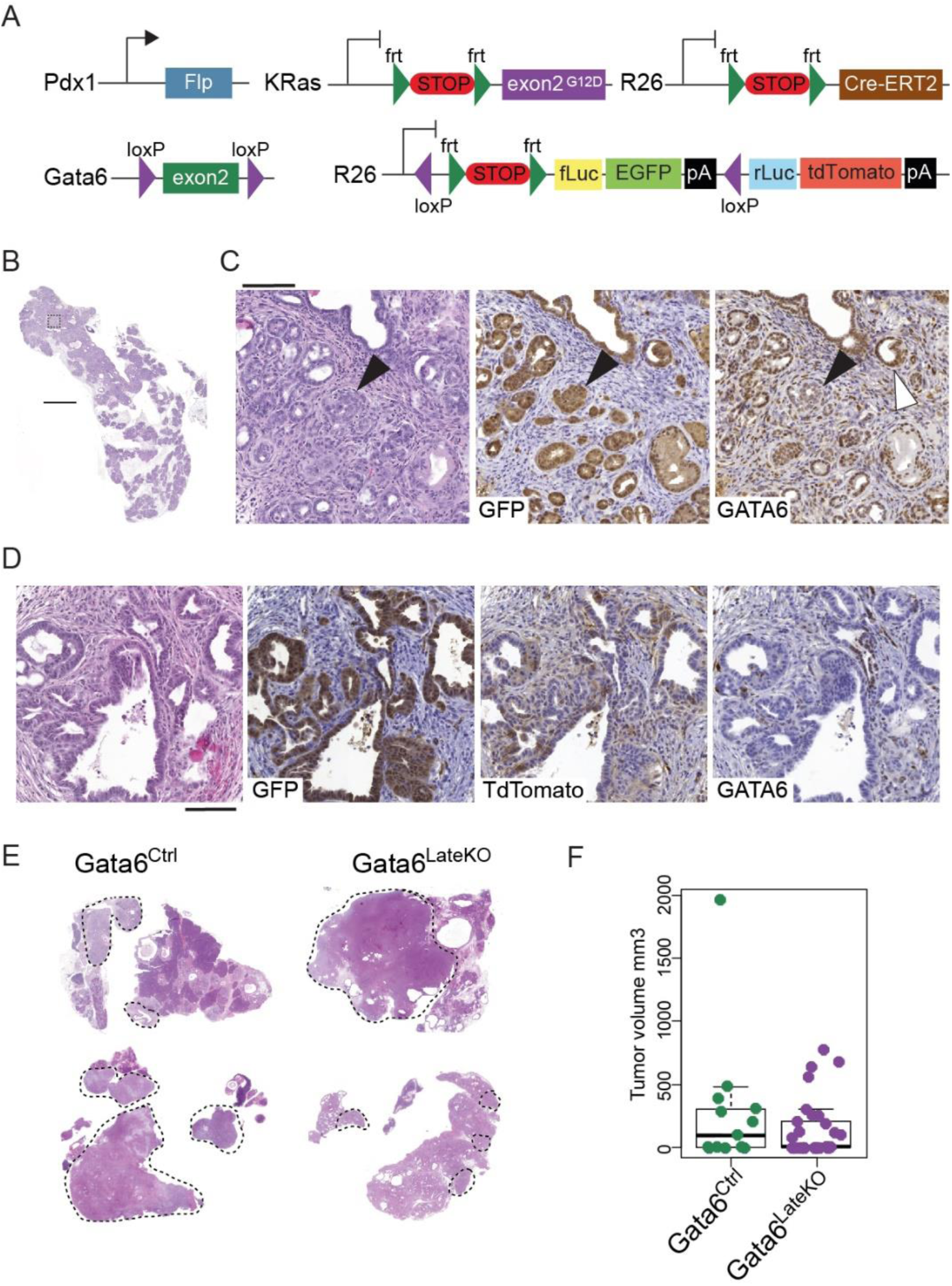
A next-generation mouse model for conditional *Gata6* deletion. A) Schematic representation of the alleles used to generate the Gata6^LateKO^ mouse model. B) Representative H&E image of the pancreas of a 20-week Pdx1-Flp and KRas^FSF-G12D^ mouse. Scale bar: 2mm. C) Images showing H&E and expression of GFP and Gata6, in a magnified region of the pancreas shown in B (dotted square). Scale bar: 100µM. D) Representative images of a Gata6^LateKO^ pancreas after TMX administration, showing H&E and GFP, TdTomato, or Gata6 expression. Scale bar: 100µM. E) Representative images of the pancreas of Gata6^Ctrl^ or Gata6^LateKO^ mice with variable tumor size. F) Quantification of the tumor volume from subsets of Gata6^Ctrl^ (n= 13) and Gata6^LateKO^ (n= 30) mice.

Flp-dependent recombination efficiency varied widely, ranging from <5% to >90% of the pancreas, and no malignant lesions were observed in pancreata having <30% of recombination, measured by IHC for the GFP reporter (not shown). We included in our analyses only mice where Flp-mediated recombination reached at least 30% of pancreatic epithelial cells. By 20 weeks of age, KFC mice developed throughout the pancreas multiple low- and high-grade PanIN lesions that stained positive for GFP (Figure 3B-C). In Gata6^wt^ mice, Gata6 was detected in the majority of epithelial cells. Occasionally, reduced Gata6 expression was observed in high-grade PanINs (Figure 3C, black arrowhead) compared to low-grade lesions (white arrowhead), suggesting that spontaneous Gata6 loss might occur at the low-high PanIN transition. We therefore administered tamoxifen (TMX) around 20 weeks of age to induce the deletion of *Gata6* in KRas^G12D^-expressing cells and generate Gata6^LateKO^ KFC mice. TMX induced efficient Cre-dependent recombination, as demonstrated by TdTomato expression in GFP-positive lesions (Figure 3D). These lesions were also consistently Gata6-negative, indicating that TdTomato is a reliable reporter of Cre-dependent recombination (Figure 3D). Importantly, no recombination was detected in Gata6^loxP/loxP^ mice not receiving TMX, as assessed by Gata6 IHC (not shown) and no GFP or TdTomato expression was detected in the stroma (Figure 3D). These observations allow excluding any relevant leakiness. Mice were sacrificed at 65 weeks or when they became moribund. From a cohort of 82 mice, 43 were Gata6^LateKO^ and 39 were controls (Gata6^Ctrl^); the latter included 28 Gata6^wt/wt^ and 3 Gata6^wt/loxP^ mice receiving TMX and 8 Gata6^loxP/loxP^ mice not receiving TMX. Gata6^Ctrl^ and Gata6^LateKO^ mice developed highly heterogeneous tumors of widely varying sizes, and no significant difference in size was observed (Figure 3E-F). The experimental design did not allow for Kaplan Maier survival analysis and we did not observe that Gata6^LateKO^ mice became moribund significantly earlier than controls (not shown). These results validated our mouse model.

### *Gata6* loss in tumors favors the basal phenotype, metastases, and lung tropism

We used Krt14 expression as a proxy for the basal phenotype in mouse PDAC, since the Tp63 staining did not give reliable results (not shown). The proportion of Krt14^pos^ tumors was significantly higher in Gata6^LateKO^ mice than in Gata6^Ctrl^ mice [25/43 (58.1%) vs. 13/39, (33.3%)] (P=0.029, Figure 4A). Importantly, among Gata6^Ctrl^ mice, all Krt14^pos^ tumors were Gata6^neg^ (Gata6^Loss^). When comparing tumors based on Gata6 expression, 38/56 (67.8%) Gata6^neg^ tumors (Gata6^LateKO^ + Gata6^Loss^) were basal, while none of the Gata6^pos^ ones was (P=7.6e-10, Figure 3A). This ultimately confirmed that GATA6 loss is necessary but not sufficient for the expression of the basal phenotype.

**Figure 4.**
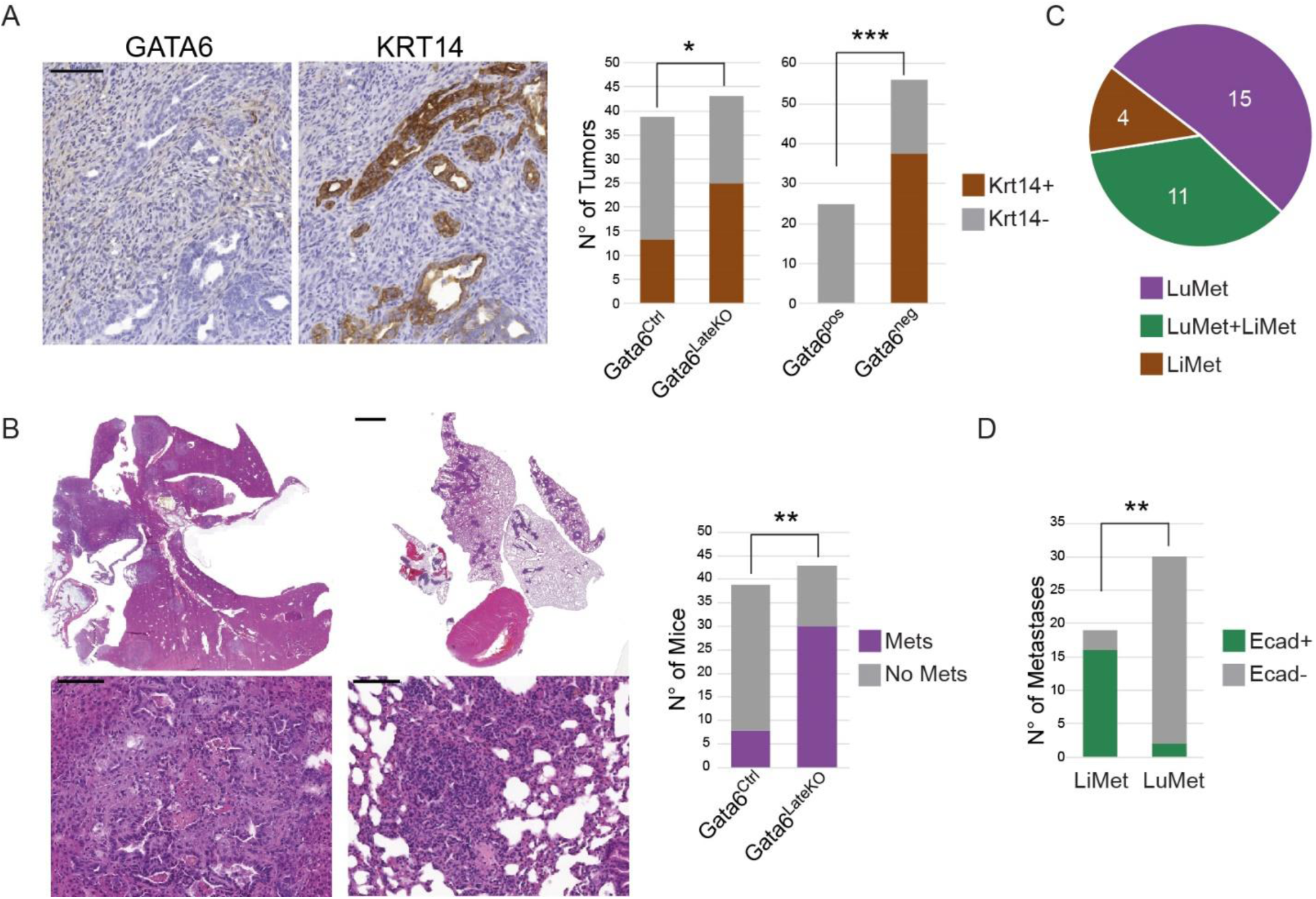
Gata6 loss in tumors leads to a basal-like phenotype and increased metastatic potential with lung-specific tropism. A) Expression of Gata6 and Krt14 in a representative Gata6^LateKO^ PDAC, detected by IHC (left) and quantification of Krt14 expression in tumors classified either by genotype (Gata6^Ctrl^ n= 39 and Gata6^LateKO^ n=43) or by Gata6 expression (Gata6^pos^ n= 26 and Gata6^neg^ n=56). *P<0.05, ***P<0.001 Scale bar: 100µm. B) Representative H&E images of liver and lung metastases in a Gata6^LateKO^ mouse and quantification of metastasis occurrence in Gata6^Ctrl^ (n= 39) and Gata6^LateKO^ (n=43) mice. **P<0.01. Scale bar: 2mm top, 100µm bottom. C) Distribution of metastases to the liver (LiMet) or to the lung (LuMet) in Gata6^LateKO^ mice. D) Quantification of E-cadherin IHC in Gata6^LateKO^ liver (n= 19) and lung (n=30) metastases, **P<0.01.

Patients with basal PDACs have worse outcome^1–4 9^. Congruently, we observed that significantly more Gata6^LateKO^ mice had clear signs of disease progression as reflected by significantly more metastases (30/43, 69.8%) than the Gata6^Ctrl^ controls (8/39, 20.5%) (P=8.5e-06, Figure 4B). Importantly, all 8 Gata6^Ctrl^ mice with metastases had Gata6^neg^ tumors (primary and metastatic). To strengthen our observations, we analyzed an independent cohort of KFC mice where transposon-based random mutagenesis was induced around week 20 in KRas^G12D^-expressing cells (KFC-SB, unpublished). Gata6 expression was lost, broadly or focally, in 19/29 KFC-SB mice (65.5%) and 18 of them (94.5%) had metastases, while only 3/10 mice with Gata6^pos^ tumors (30%) had metastases (P= 0.001, Supplementary Figure 5A). These data indicate that Gata6 is an efficient suppressor of metastasis in murine KRas^G12D^-driven PDAC.

Among the Gata6^LateKO^ mice, 15/30 (50%) had only lung metastases, 4/30 (13.3%) had only liver metastases, and 11/30 (36.7%) had both (Figure 3C). This result differs from findings in patients, where the liver is the most common site of metastases^19^. There is evidence that tumor cells are heterogeneous and must, in addition, be highly plastic to form metastases^20 21^. This degree of plasticity influences the organotropism of PDAC metastatic cells, whereby cells that cannot fully revert the epithelial phenotype colonize preferentially the lungs^22^. Among Gata6^LateKO^ mice, liver metastases were significantly more often E-cadherin^pos^ than lung metastases as detected by IHC (16/19, 84.2% E-cadherin^pos^ LiMet; 2/30, 6.7% E-cadherin^pos^ LuMet, P= 3.69e-08) (Figure 4D, Supplementary Figure 5B). This observation is consistent with data showing undetectable E-cadherin in primary cells derived from a lung metastasis^23^ and might indicate that Gata6^LateKO^ cells are more limited in their ability to efficiently reactivate the epithelial program.

### Gata6^LateKO^ primary tumor cells are more proliferative and chemoresistant

We successfully established primary cell lines from Gata6^pos^ (n=5), Gata6^LateKO^ (n=15) and Gata6^Loss^ (n=6) tumors. While Gata6^pos^ and Gata6^LateKO^ cell lines were homogeneously positive and negative for Gata6, respectively, Gata6^Loss^ lines displayed a more heterogeneous expression pattern (Figure 5A). The majority (10/15) of Gata6^LateKO^ and 4/6 Gata6^Loss^ lines were positive for Krt14 while 5/5 Gata6^pos^ were negative (Supplementary Figure 6A). Additionally, ΔNp63 mRNA was significantly higher in Gata6^LateKO^ and Gata6^Loss^ cells (Figure 5B), accompanied by a similar trend in a set of basal markers (Runx3, S100a2, and Krt14) but not classical/progenitor markers (Pdx1 and Hnf4a, Supplementary Figure 7A) indicating that these cells preserve basal features *in vitro*. Gata6^LateKO^ and Gata6^Loss^ cells were significantly more proliferative compared with Gata6^pos^ cells (Figure 5C). The migratory capacity of KFC cells was highly variable and no statistical differences were observed, although some of the Gata6^LateKO^ and Gata6^Loss^ cells showed high migratory potential (Supplementary Figure 6B). No difference was observed in the invasive capacity *in vitro* (Supplementary Figure 4C).

**Figure 5.**
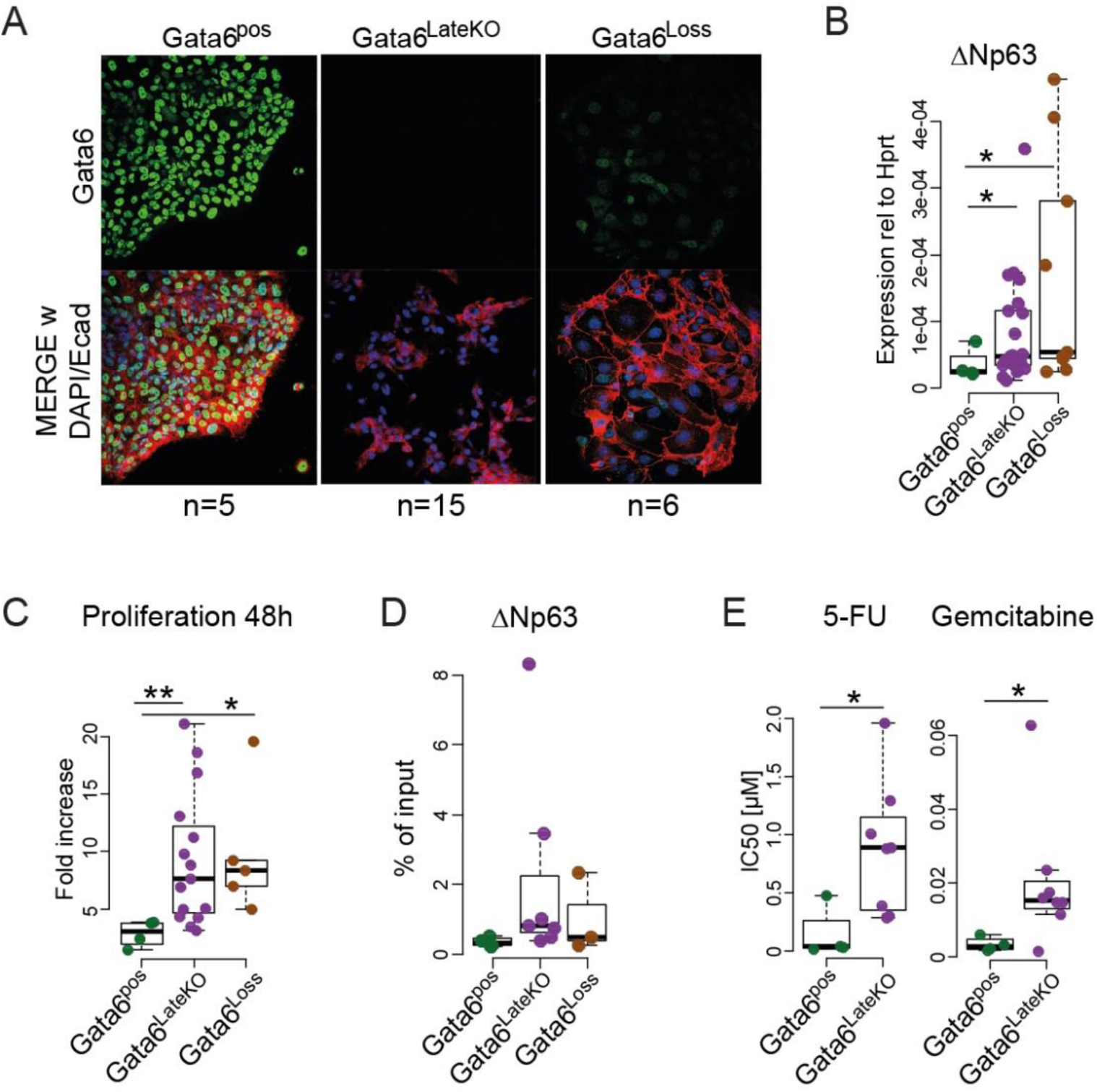
Primary cells from Gata6^LateKO^ tumors are more proliferative and chemo-resistant *in vitro*. A) Representative immunofluorescence images of primary KFC tumor cells isolated from Gata6^pos^, Gata6^LateKO^, and Gata6^Loss^ mice. Top: expression of Gata6 (green). Bottom: merged Gata6 (green), E-cadherin (red) and DAPI (blue) stainings. B) Expression of ΔNp63 measured by RT-qPCR. C) Proliferation of primary KFC cells from the indicated groups, represented as fold increase in cell number 48 h after seeding. Gata6^pos^ n=4, Gata6^LateKO^ n=15, Gata6^Loss^ n=5. D) H3K27ac binding to the promoter of ΔNp63, detected by ChIP-qPCR in primary KFC cells. Data are represented as % of input chromatin. Gata6^pos^ n=4, Gata6^LateKO^ n=7, Gata6^Loss^ n=3. E) Graphs representing the IC50 values measured for primary KFC cells upon treatment with 5-FU and Gemcitabine in cytotoxicity assays. Gata6^pos^ n=4, Gata6^LateKO^ n=8. B-E: each dot represents the average value of at least three independent experiments for each tumor cell line. *P<0.05, **P<0.01.

Next, we investigated the epigenetic contributions to our observations, since this mechanism has been shown to control the emergence of the basal phenotype in PDAC cells^11 13 16^. ChIP-qPCR for H3K27ac, a marker of open chromatin, showed higher enrichment on the *ΔNp63* promoter in *Gata6^LateKO^* and *Gata6^Loss^* cells (Figure 5D). No significant differences were observed for the promoters of a subset of basal (*Runx3*, *S100a2*, *Krt14*) and classical (*Pdx1* and *Hnf4a*) genes (Supplementary Figure 7B). Hence, based on this analysis, *Gata6* deletion does not appear to cause widespread remodeling of chromatin accessibility in KFC cells.

Lastly, we evaluated the relationship between GATA6 status and the response of tumor cells to chemotherapeutic agents commonly used for the treatment of PDAC. Patients with low GATA6^low^ or basal-like PDAC respond worse to 5-FU-based adjuvant treatments^9 10^. Consistently, Gata6^LateKO^ cell lines were significantly more resistant to 5-FU than Gata6^pos^ cells (Figure 5E). In contrast to the findings in patients, however, Gata6^LateKO^ cells were also more resistant to gemcitabine (Figure 5E). These observations indicate that the KFC cell line panel generated in this study recapitulates some features of the human disease, including the high inter-patient heterogeneity.

### GATA6 loss favors cell plasticity and immune escape

To pinpoint the mechanism underlying Gata6 basal- and metastasis-suppressive function, we performed RNAseq analysis of Gata6^pos^, Gata6^LateKO^ and Gata6^Loss^ primary tumor cells. A recent multi-omics analysis of primary mouse PDAC cells identified two transcriptomics-based clusters: C1 (Mes-C1) and C2 (Epi-C2, including 3 subclusters C2a, C2b, and C2c) roughly overlapping with the basal and classical subtypes in patients, respectively^24^. Reference cell lines from that analysis were included in our experiment^24^. KFC cells could be assigned to three clusters: C1, C2a, and C2b/c. All but one Gata6^pos^ lines fell into cluster Epi-C2b/c and one was assigned to the Epi-C2a cluster. In contrast, all lines assigned to the Mes-C1 cluster were either Gata6^LateKO^ or Gata6^Loss^ cells, supporting the strong anti-EMT role of GATA6 (Figure 6A, Supplementary Figure 8A). Eight of the Gata6^LateKO^ cell lines included in the RNAseq analysis were isolated from primary tumors that had metastasized. Interestingly, the Mes-C1 cluster included only cells from lung-tropic tumors, while the Epi-C2b/c cluster only included cells from tumors that also generated liver metastases (Figure 6A). This data further supports that PDAC cells with a strong mesenchymal phenotype colonize preferentially the lungs.

**Figure 6.**
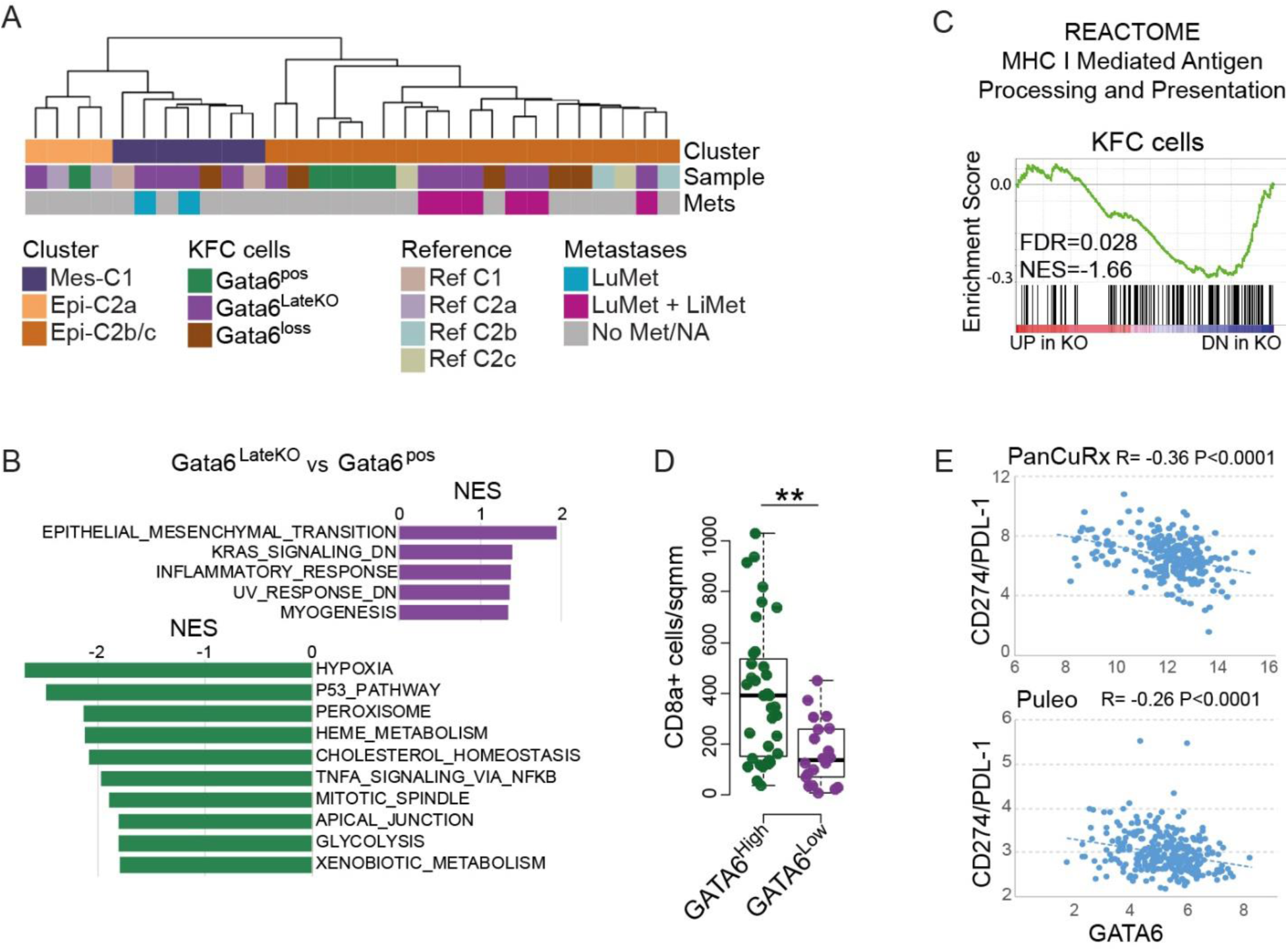
GATA6 loss favors cell plasticity and immune escape. A) Hierarchical clustering of KFC cells and reference cell lines according to RNAseq analysis. Gata6^pos^, Gata6^LateKO^ and Gata6^Loss^ primary cells were assigned to the described clusters Mes-C1 (mesenchymal), Epi-C2a (epithelial), and Epi-C2b/c (epithelial). B) Gene set enrichment analysis of genes differentially regulated between Gata6^LateKO^ and Gata6^pos^ cells. C) Enrichment of the gene set “MHCI mediated antigen processing and presentation” in KFC cells. D) Quantification of CD8α-positive T cells in GATA6^High^ and GATA6^Low^ patient-derived tumors, detected by IHC. *P<0.05. E) Correlation between GATA6 and CD274 (coding for PDL-1) mRNA levels in the indicated datasets.

Subsequently, we performed gene set enrichment analysis (GSEA) with all possible comparisons. We found “EMT” as the most highly enriched gene set among the genes upregulated in Gata6^LateKO^ or Gata6^Loss^ cells compared with Gata6^pos^. On the other hand, an “apical junctions” gene set was enriched among the genes upregulated in Gata6^pos^ vs Gata6^LateKO^ cells (Figure 6B). These results reveal the existence of complementary effects, primarily supporting the acquisition of more mesenchymal features, which are characteristic of metastatic PDAC cells. GSEA showed additional similarities between the Gata6^LateKO^ and the Gata6^Loss^ cells, when compared to the Gata6^pos^ cells, including the enrichment of the “KRAS signaling_DN” gene set among the up-regulated genes. In contrast, the “Hypoxia”, “p53-pathway”, and “metabolism-related” gene sets were down-regulated (Figure 6B, Supplementary Figure 8B, Supplementary Table 1). Moreover, we also observed significant differences between Gata6^LateKO^ and Gata6^Loss^ cells (Supplementary Figure 8B,C). Finally, when comparing Gata6^LateKO^ cells with epithelial or mesenchymal features, EMT was clearly up-regulated in Mes-Gata6^LateKO^, while two “MYC targets” gene sets, cell cycle-related gene sets, and DNA-repair-related ones were upregulated in Epi-Gata6^LateKO^ (Supplementary Figure 8D), confirming that Gata6^LateKO^ cells are diverse. Therefore, heterogeneity in pancreatic cells is a defining characteristic not only of human but also of genetically engineered mice used to model this disease.

Interestingly, the MHC class I genes *H2-d1* and *H2-k1* and the immunoproteasome gene *Psmd8* were among the most significantly down-regulated genes in the Gata6^LateKO^ cells, suggesting that Gata6 loss might induce immune escape, thereby supporting higher metastatic potential. Congruently, the gene set “MHC I Mediated Antigen Processing and Presentation” was significantly enriched among genes down-regulated in the Gata6^LateKO^ cells (Figure 6C). IHC analysis of GATA6^low^ patient tumors revealed a significantly decreased infiltration of CD8α+ T cells compared to GATA6^high^ tumors (Figure 6D). To expand these observations, we explored the available patient-derived datasets for evidence of GATA6 involvement in immune escape. The Puleo cohort showed the most consistent results: MHC I-mediated antigen processing and presentation and the estimated abundance of CD8+ T cells^25^ were significantly lower in GATA6^low^ tumors (Supplementary Figure 9A,B). Furthermore, the T cell checkpoint activator PDL-1 (encoded by the *CD274* gene) was negatively correlated with GATA6 expression in 3/5 datasets (Figure 6E and Supplementary Figure 9C). Interestingly, *CD274* and several genes related with antigen processing and presentation (*PSMD8*, *PSMD9*, *B2M*) had GATA6 peaks on the promoter in the ChIP-Seq we performed in PDAC cells^9^, suggesting that GATA6 might directly regulate a subset of them. Taken together, our data suggest that GATA6 loss in PDAC can facilitate immune escape, favoring metastasis.

## DISCUSSION

The recent -omics technological revolution already led to major leaps forward in our understanding of PDAC biology. In particular, transcriptomic-based tumor taxonomy revealed important differences and commonalities among PDACs. A detailed understanding of the molecular events driving the different phenotypes, particularly the highly aggressive basal program, will increase the chance of a successful translation of basic knowledge into clinical intervention.

Here we show that loss of GATA6, a major regulator of epithelial identity, is necessary, but not sufficient, for the basal phenotype in patient-derived samples and in a next-generation mouse model in which *Gata6* deletion was induced at the time of KRas^G12D^-driven high-grade PanIN formation (Gata6^LateKO^).

Multiple lines of evidence link GATA6 loss to the basal phenotype^1 2 9 10^. Our data from the Gata6^LateKO^ mice ultimately identify GATA6 as a molecular gatekeeper restricting cell plasticity to maintain lineage-specific programs in PDAC. Gata6 loss is necessary to allow the expression of the basal program, but additional downstream or parallel events are required. Analysis of patient-derived samples revealed that HNF1A and HNF4A might act as further molecular barriers to maintain the classical gene program when GATA6 is lost, possibly through the regulation of a shared subset of genes. GATA6, HNF1A, and HNF4A are all involved in cell fate determination of the pancreatic lineage during development and are highly expressed in the classical/progenitor PDACs^1 26–29^. Non-basal GATA6^low^ tumors in patients displayed intermediate levels of GATA6 expression, suggesting the existence of a threshold below which HNF1A and HNF4A expression is lost and the full basal program is established. Such a threshold does not exist in mouse tumors in our model, where Gata6 is lost at the genomic level. Therefore, loss of HNFs expression is likely the result of multiple regulatory events. Lomberk and colleagues described that GATA6 influences the expression of HNFs to maintain the classical phenotype in patient-derived xenografts^14^. Our work confirms this hierarchy, whereby GATA6 acts upstream of both HNF1A and HNF4A to block the basal program (Figure 2E). Importantly, *HNF4A* was epigenetically silenced in GATA6^low^/Basal PDX-derived cells, suggesting that epigenomic remodeling is a crucial event downstream of GATA6 loss, to allow the full classical-basal switch.

The relationship between GATA6 and ΔNp63 in controlling the classical and basal phenotypes is complex. Our data and the reanalysis of available datasets suggest that neither GATA6 loss nor ΔNp63 expression are sufficient for a full phenotype switch, but both events are necessary and there is evidence of a cross-regulation whereby GATA6 downregulation allows for expression of ΔNP63, which in turn contributes to keep GATA6 inhibited. Similar regulatory relationships might be true for HNF4A and HNF1A.

While the *in vivo* findings and the correlations observed in patient-derived samples were highly consistent, *in vitro* modulation of GATA6 and ΔNp63 expression in cell lines yielded variable results suggesting highly context-dependent effects including roles for the stroma and the immune system. In particular, BxPC3 are KRAS wt, thus representing a rare subset of PDAC patients. The different behavior we observed in BxPC3 and L3.6pl cells after GATA6 overexpression might reflect the contribution of mutant KRAS. These differences might indicate that BxPC3 and L3.6pl cell lines do not faithfully represent the complexity of the basal-like PDACs. Indeed, only 3/6 patient-derived xenografts (PDX) originally defined as basal-like^14^ shared the H3K27ac pattern of BxPC3 and L3.6pl^11^. Our panel of primary mouse tumor cell lines represents a valuable tool for understanding the basal phenotype, adding to the PDX-derived cells described previously^14^.

We additionally show that Gata6 loss dramatically increases the rate of metastasis, thus providing a molecular link between the basal program and the metastatic potential of PDAC. A recently published transcriptomic dataset of PDAC encompassing all-stages confirmed that the basal phenotype is highly enriched among metastatic tumors^5^. A thorough characterization of primary cell lines isolated from mouse tumors revealed that Gata6 loss results in higher plasticity and possibly immune evasion, both characteristics of metastatic cells.

Cellular plasticity is one hallmark of metastatic cells. While EMT is required to initiate metastatic spread, the reverse process - MET - is necessary for the growth of metastases at distant sites and cells that cannot revert the EMT are not able to grow metastases in mouse models^20 21^. The degree of plasticity seems to play an important role in defining the mode of dissemination^30^ and the organotropism of metastases, with more epithelial-like cells forming metastases preferentially the liver and more mesenchymal-like cells favoring lung metastases^22^. Gata6^LateKO^ mice preferentially developed lung metastases, mostly E-cadherin-negative, indicating that Gata6-KO cells might not be able to revert to a fully differentiated status. This is consistent with the crucial role of GATA6 in establishing epithelial and pancreatic cell identity^7^.

Disseminating tumor cells must overcome multiple hurdles during their path to the metastatic site, among them the immune surveillance. Suppression of antigen processing and presentation is one mechanism of immune evasion that tumor cells have hijacked from viruses^31 32^. We observed that GATA6 loss in tumors from patients and mice decreased the expression of the antigen processing and presentation machinery and that infiltration of CD8-positive T cells was reduced in GATA6^low^ tumors in patients, possibly indicating a more efficient immune evasion. Accordingly, Gata6 knock-out favored T cell-mediated tumor cell killing in an *in vivo* CRISPR screening^31^, suggesting that the immunogenicity gene program is embedded within the GATA6-dependent epithelial cell identity program. We reported similar findings for another master regulator of the epithelial cell identity, NR5A2, which actively inhibits an inflammatory gene expression program in the normal pancreas^33^. The polycomb repressive complex 2 (PRC) was recently shown to silence the MHC class I antigen presentation pathway^32^; interestingly, we observed GATA6 peaks on the promoter of EZH2 and EED in our published ChIP-Seq, and their transcripts were mildly down-regulated in RNA-Seq upon GATA6 silencing^9^. This points to an indirect role of GATA6 in modulating the antigen processing and presentation machinery, possibly through PRC2.

In summary, we show here that a GATA6-centered gene regulatory network functions as a gatekeeper of cell identity and blocks cell plasticity and immune evasion, thus providing a molecular link between the basal-like phenotype and metastasis. The Gata6^LateKO^ mouse model is therefore a valuable preclinical tool to study the most aggressive subtype of PDAC.

## Acknowledgements

Work in the lab of PM was supported by the grant P27361-B23 from the Austrian Science Grant (FWF) and by contributions from the Fellinger Krebsforschung foundation and the Ingrid Shaker-Nessmann Krebsforschungsvereinigung foundation. Patients were not involved in the design of this study. RU and GL were supported by The Linda T. and John A. Mellowes Endowed Innovation and Discovery Fund, the Theodore W. Batterman Family Foundation, Inc and NIH Grants: R01 DK52913 and R01 CA178627. Work in the lab of F.X.R. was supported, in part, by grant RTI2018-101071-B-I00 (Ministerio de Ciencia, Innovación y Universidades, Madrid, Spain). CNIO is supported by Ministerio de Ciencia, Innovación y Universidades as a Centro de Excelencia Severo Ochoa SEV-2015-0510. The authors declare no conflict of interest.

## Authors’ contribution

PM designed the study and secured funding, analyzed the experiments and prepared the manuscript. BK performed most of the experiments and participated in writing the manuscript. NH and VI managed the mouse colony and performed experiments. JM performed some bioinformatics analyses. RO, SM, and RR performed and analyzed the RNA-Seq experiment. HPD and JS did the pathology assessments. MS and EG provided access to resection tissues. GL and RU performed the analysis of epigenomic data from PDX-derived cells. BS and DS provided the dual recombinase mouse strain. FXR provided the Gata6 flox mice and participated in the experimental design and manuscript preparation.

## Supplementary Material

### Methods

### Mice

The following mouse strains were described previously: *FSF-Kras^G12D/+^, Pdx1-Flp*, *FSF-R26^CAG-CreERT2/+ 17^*, *Gata6^loxP/loxP16^*,, T2/Onc^34^, Rosa26-LSL-SB13^35^, and *Rosa26-CAG-loxP-frt-Stop-frt-FireflyLuc-EGFP-loxP-RenillaLuc-tdTomato* (referred to as R26^Dual^)^18^. The strains were interbred to obtain mice expressing oncogenic KRasG12D in the pancreas, together with the other alleles as mentioned in the text. Flp-dependent recombination was verified by PCR and GFP expression, and mice with no recombination were excluded from the analyses.

Male and female mice were given Tamoxifen-containing diet (Cre-active-400, Genobios) or 4-OHT intraperitoneal injections (10 mg/mL in corn oil, Sigma, 2 mg injected / animal) between 20 and 30 weeks of age to induce Cre activity. Organs were collected at 65 weeks, or earlier if mice became moribund.

Mice were bred and maintained at the Medical University of Vienna, under pathogen-free conditions. Animals had unlimited access to standard food and water and a light-dark cycle of 12 h at 22 °C temperature, in accordance to guidelines of the institute and federal regulations. All experiments were conducted in compliance with Animal Ethics Committee of the Medical University of Vienna and federal laws. Ethical approval was obtained for all experiments (BMWFW-66.009/116-WF/V/3b/2015)

#### Patient cohort, resected PDAC samples and data

Sixty patients with a diagnosis of PDAC undergoing surgery with curative intent between 2015 and 2016 at the Department of Surgery, Medical University of Vienna, were retrospectively defined as study cohort and the corresponding FFPE tumor samples were prepared for further analysis. Patient and tumor characteristics were collected from the institutional database. The study was approved by the local ethics committee of the Medical University of Vienna (“Ethikkommission”, protocol no. 1753/2014).

#### Patient-derived expression datasets

The following datasets were downloaded from the indicated source and used in the analyses: Collisson (GSE17891, Array-based), Moffitt (GSE71729, Array-based), Bailey (Supplemental table from the publication, RNA-Seq), TCGA (UCSC-XENA browser, RNA-Seq), Puleo (E-MTAB-6134, Array-based), PanCuRx (EGAS00001002543, RNA-Seq).

#### Histopathology and immunohistochemistry (IHC)

Tissues were fixed in phosphate buffered formalin and embedded in paraffin. For histopathological analysis, tissues were serially sectioned (3 µm). Sections were stained with hematoxylin and eosin according to standard protocols. Immunostaining for GATA6, KRT14, GFP, TP63 and E-cadherin were performed according to standard protocols with DAKO reagents. DAB was used as chromogen, nuclei were counterstained using hematoxylin. Stained sections were scanned using 3DHISTECH Pannoramic MIDI Slidescanner and analyzed using Pannoramic Viewer 1.15.4 Software. The antibodies used are indicated in Supplementary Table 1.

#### Cell lines

Primary cell lines were established from KFC tumors and metastases. Tissue was finely minced, digested in collagenase IV (Thermo Fisher), resuspended 1:1 in Matrigel (Sigma), seeded in 50µl droplets and cultivated in RPMI-1640 supplemented with 10% FBS and Penicillin/Streptomycin (Sigma). Successfully isolated cell lines were cultured in RPMI-1640 supplemented with 10% FBS and Penicillin/Streptomycin (Sigma) under standard 2D conditions.

HEK293T and PaTu8988S cells were cultured in DMEM supplemented with 10% FBS. L3.6pl and BxPC3 cells were cultured in RPMI-1640 supplemented with 10% FBS under standard conditions (37°C, 5% CO_2_, 20% O_2_). All the established cell lines were already available in the laboratory and were periodically tested for mycoplasma contamination by PCR.

#### Cytotoxicity assay

Cells were seeded at low density (2000 cells/well) in a 96-well plate and treated as previously described^9^. After 72 h, cell viability was measured by an MTT-based assay (Biomedica) according to the manufacturer’s instructions.

#### Matrigel invasion assay

Transwells (Falcon, 8 µm) were coated with Matrigel (Sigma) diluted 1:10 in sterile PBS. Cells (10^5^) were seeded onto Matrigel in serum-free RPMI-1640 and were allowed to invade towards RPMI-1640 containing 10% FBS added to the lower compartment. Cells that invaded the matrigel and were attached to the lower side of the transwell membrane were fixed after 24 h using 4% PFA, stained with DAPI and counted using a fluorescent microscope and the Definiens software.

##### Wound-healing assay

Cells were grown until they reached confluence, then the monolayer was scratched using a sterile 10µl pipette tip. Cells were washed once with PBS to remove floating cells and then changed to medium containing 1% FBS. The closure of the wound was monitored using a microscope with automatic image capture function (Zeiss).

##### Proliferation assay

Cells (2x10^4^) were seeded in triplicates in a 6-well plate. Each day, for four consecutive days, cells were detached using trypsin and counted by CASY Cell Counter.

##### Plasmids, transfection and infection

Lentiviral vectors expressing non-targeting and GATA6-targeting shRNAs were purchased from SIGMA-Aldrich (MISSION sh-RNA). GATA6 cDNA was cloned into the GFP-expressing FG12 lentiviral vector for overexpression in PDAC cells^9^. Virus-packaging HEK293T cells were transfected with standard calcium phosphate protocol, supernatant was collected 48h after transfection, filtered, and used to infect PDAC cells. Successfully infected cells were selected with puromycin or with FACS-sorting for GFP.

##### Protein analysis

Total protein lysates were obtained from cell culture dishes using Laemmli-Buffer. Sonicated samples were used for SDS-PAGE-western blotting using standard protocols. The antibodies used are indicated in Supplementary Table 2.

##### Immunofluorescence

Cells were seeded onto sterile glass cover slips, fixed with 4% PFA and permeabilized using TritonX according to standard protocols. Alexa-conjugated secondary antibodies (Life Technologies) were used, nuclei were stained with DAPI.

##### ChIP

ChIP was performed as described previously^9^. Briefly, cells were cross-linked with 1% formaldehyde for 8 minutes, and nuclei were enriched and lysed with SDS-containing buffer. Chromatin was sonicated using a Diagenode Bioruptor® instrument (15 cycles, 30 sec burst per cycle), and protein-DNA complexes were immunoprecipitated using anti-Histone H3-acetyl K27 antibody (Abcam) and A/G-agarose beads (Santa Cruz). After de-crosslinking, DNA was extracted with standard phenol-chloroform procedure and analyzed using qPCR. Primer sequences are listed in Supplementary Table 2.

##### Gene expression analysis

Total RNA was extracted from cell culture dishes using Trizol followed by phenolchlorofom extraction. RNA was retrotranscribed with LunaScript RT mix (NEB) following manufacturer’s instructions. qPCR was performed on cDNA using SYBR-green reagents and Bio-Rad CFX real time PCR instruments. Primer sequences are listed in Supplementary Table 2.

##### RNA-sequencing

Library preparation for bulk 3’-sequencing of poly(A)-RNA was done as described previously^36^. Barcoded cDNA of each sample was generated with a Maxima RT polymerase (Thermo Fisher) using oligo-dT primer containing barcodes, unique molecular identifiers (UMIs) and an adapter. 5’ ends of the cDNAs were extended by a template switch oligo (TSO) and after pooling of all samples full-length cDNA was amplified with primers binding to the TSO-site and the adapter. cDNA was tagmented with the Nextera XT kit (Illumina) and 3’-end-fragments finally amplified using primers with Illumina P5 and P7 overhangs. In comparison to Parekh et al. the P5 and P7 sites were exchanged to allow sequencing of the cDNA in read1 and barcodes and UMIs in read2 to achieve a better cluster recognition. The library was sequenced on a NextSeq 500 (Illumina) with 65 cycles for the cDNA in read1 and 16 cycles for the barcodes and UMIs in read2.

Data was processed using the published Drop-seq pipeline (v1.0) to generate sample- and gene-wise UMI tables^37^. Reference genome (GRCm38) was used for alignment. Transcript and gene definitions were used according to the ENSEMBL annotation release 75. Raw data were deposited at ENA.

##### Gene Set Enrichment Analyses (GSEA)

Differential gene expression was computed using the Comparative Marker Selection module of Genepattern, by performing pairwise comparisons. GSEA was then calculated on the ranked list of makers using the GSEA Preranked module and interrogating the Hallmarks and the C3-TFT Transcription Factor Targets gene set collections, or selected gene sets from the MSigDb database of the Broad Institute. GATA6^low^ and GATA6^high^ tumors were defined as the bottom and top quartile of patients for GATA6 expression, respectively, in all the patient-derived datasets analyzed. FDR <0.05 was considered significant.

##### Statistical analysis

Data are provided as mean±SEM. Statistical analysis was performed with VassarStat.net, R Studio, and Graph Pad Prism, using two tailed Student’s T-test for pairwise comparisons, Fisher Exact Probability test, or Pearson’s correlation test. Significance was considered for two-sided P<0.05.

**Supplementary Figure 1.**
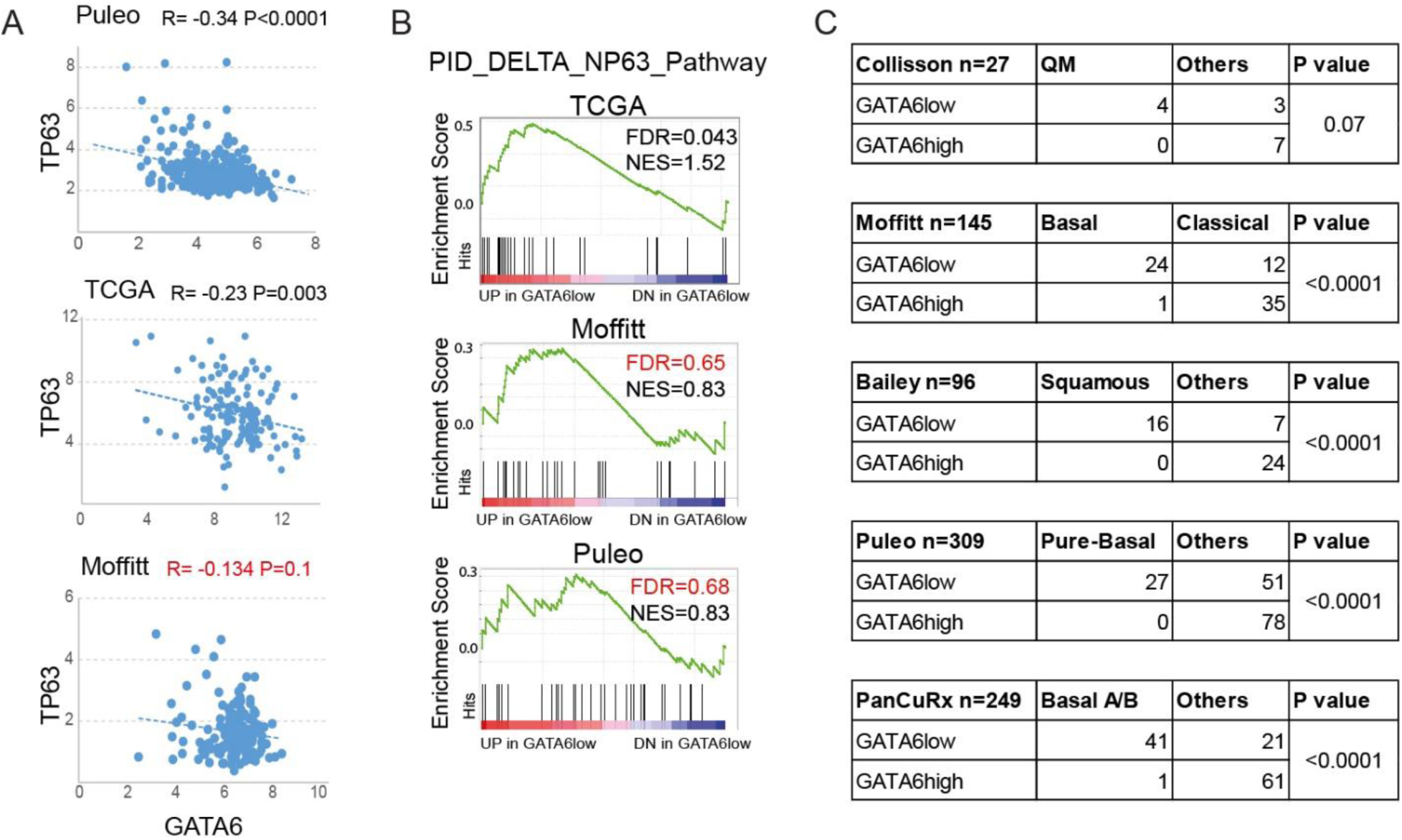
GATA6 loss in PDAC correlates with the basal phenotype. A) Correlation between GATA6 and TP63 expression in patient-derived transcriptomic datasets B) Enrichment of the gene set “ΔNp63 target genes” among the genes up-regulated in GATA6^low^ versus GATA6^high^ tumors in the indicated datasets. C) Frequency of basal and non-basal subtypes among the GATA6^low^ and GATA6^high^ tumors (bottom and top quartile, respectively) of the five datasets.

**Supplementary Figure 2.**
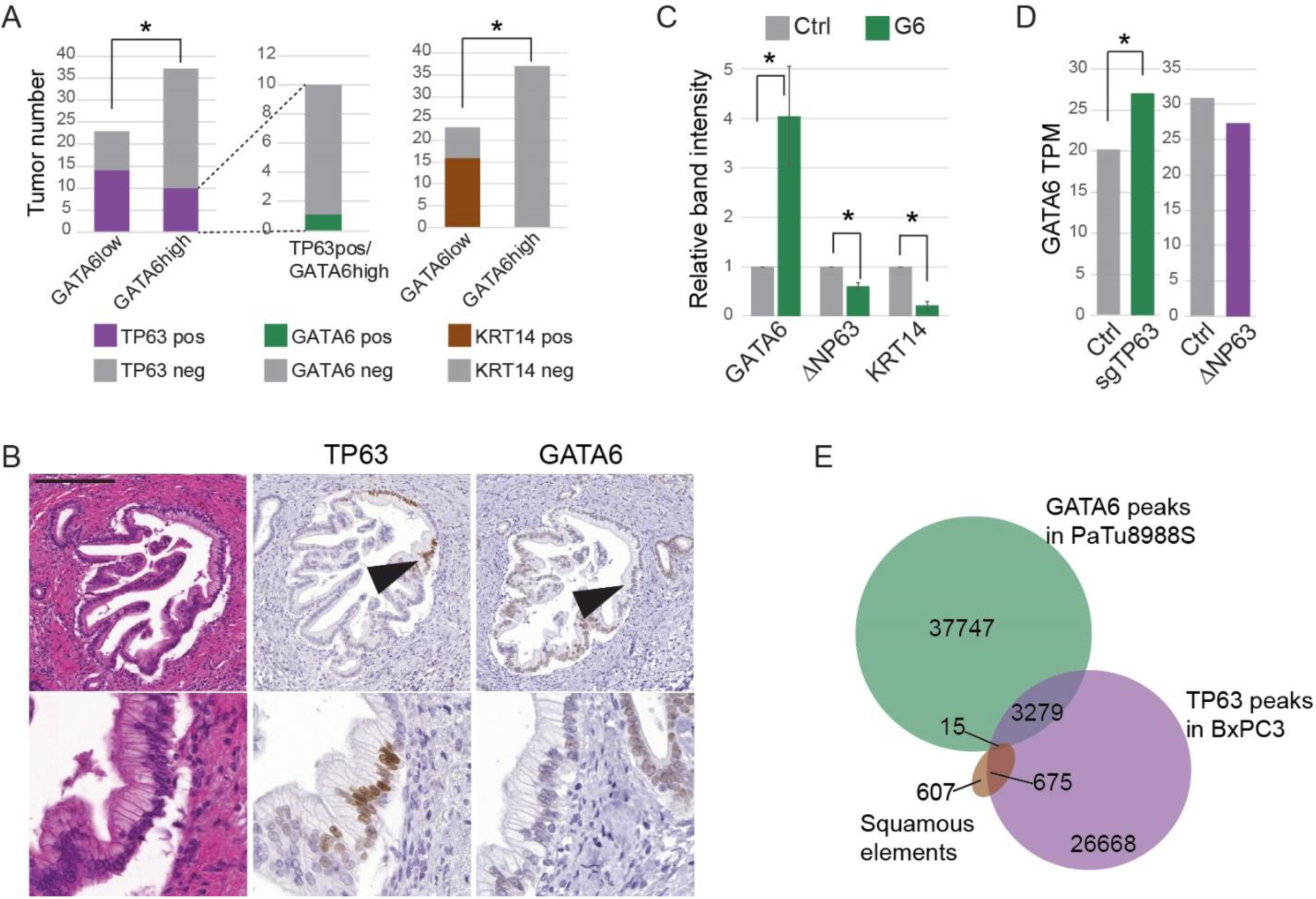
GATA6 inhibits the basal phenotype in PDAC. A) Quantification of TP63 expression in GATA6^high^ and GATA6^low^ tumors. Refers to Figure 1D. B) Representative H&E and expression of TP63 and GATA6 in a metaplastic lesion detected by IHC. Magnification of a detail indicated by the black arrowhead is shown on the bottom. Scale bar: 200µM. C) Quantification of western blot analysis depicted in Figure 1E. D) Expression of GATA6 in BxPC3 cells after CRISPR/Cas9 knock-out of TP63 (left) and in PaTu8988S cells after overexpression of ΔNp63 (right) measured by RNA-Seq (data from Somerville et al.). E) Overlap between GATA6 ChIP-Seq peaks in PaTu8988S, TP63 peaks in BxPC3, and the squamous elements defined by Somerville et al.

**Supplementary Figure 3.**
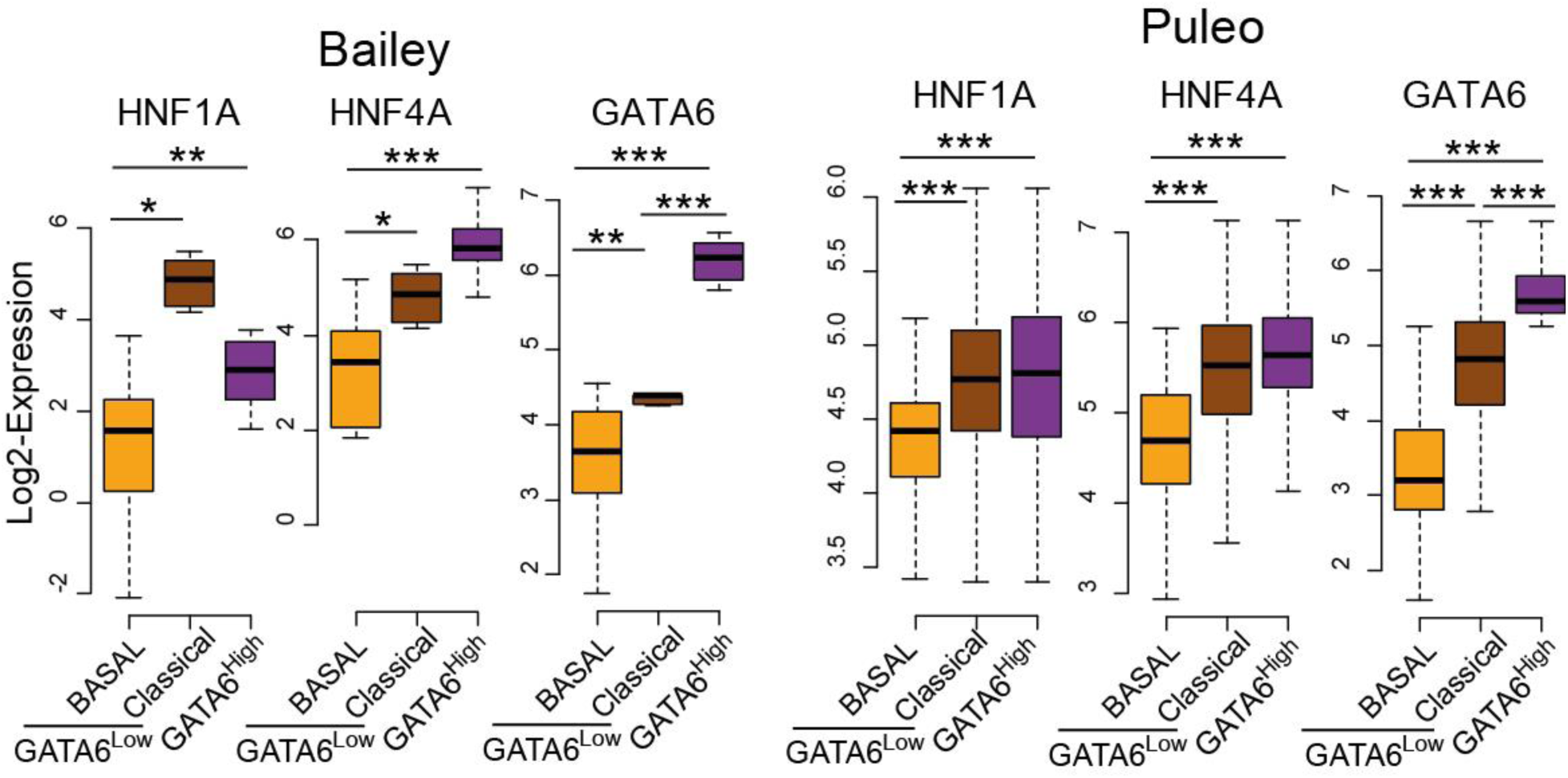
GATA6^low^ classical tumors express higher HNF1A and HNF4A. Expression of HNF1A, HNF4A, and GATA6 in the different groups of patients in the indicated datasets. *P<0.05, **P<0.001, ***P<0.0001.

**Supplementary Figure 4.**
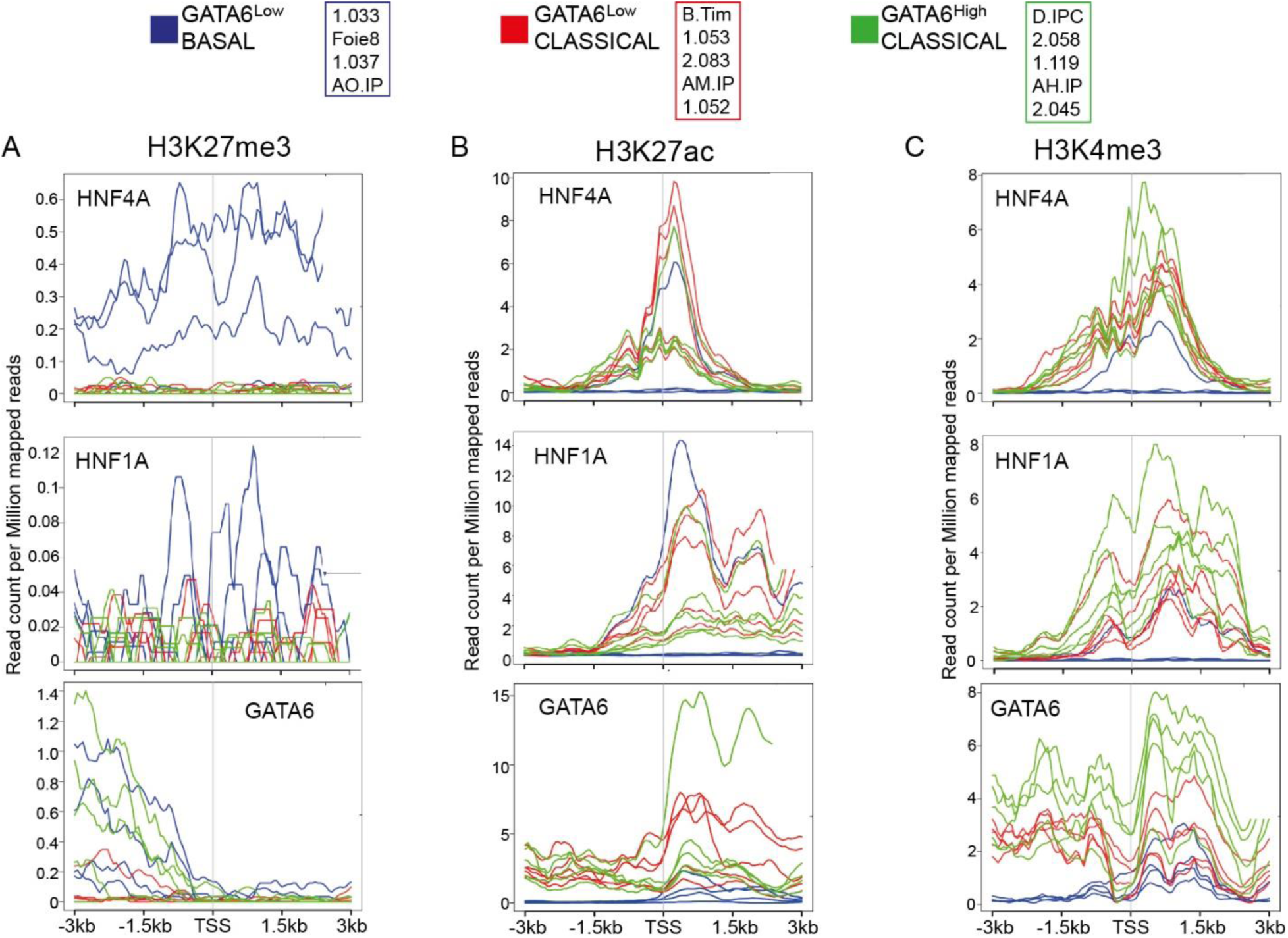
Epigenetic marks around *HNF4A*, *HNF1A*, and *GATA6* TSS differ between PDX-derived cells. A-C) Read count profile from H3K27me (A), H3K27ac (B), and H3K4me3 (C) ChIP-Seq on PDX-derived cells around the transcription start site (TSS) of *HNF4A*, *HNF1A*, and *GATA6*. Each line represents one of the cell lines indicated on the bottom.

**Supplementary Figure 5.**
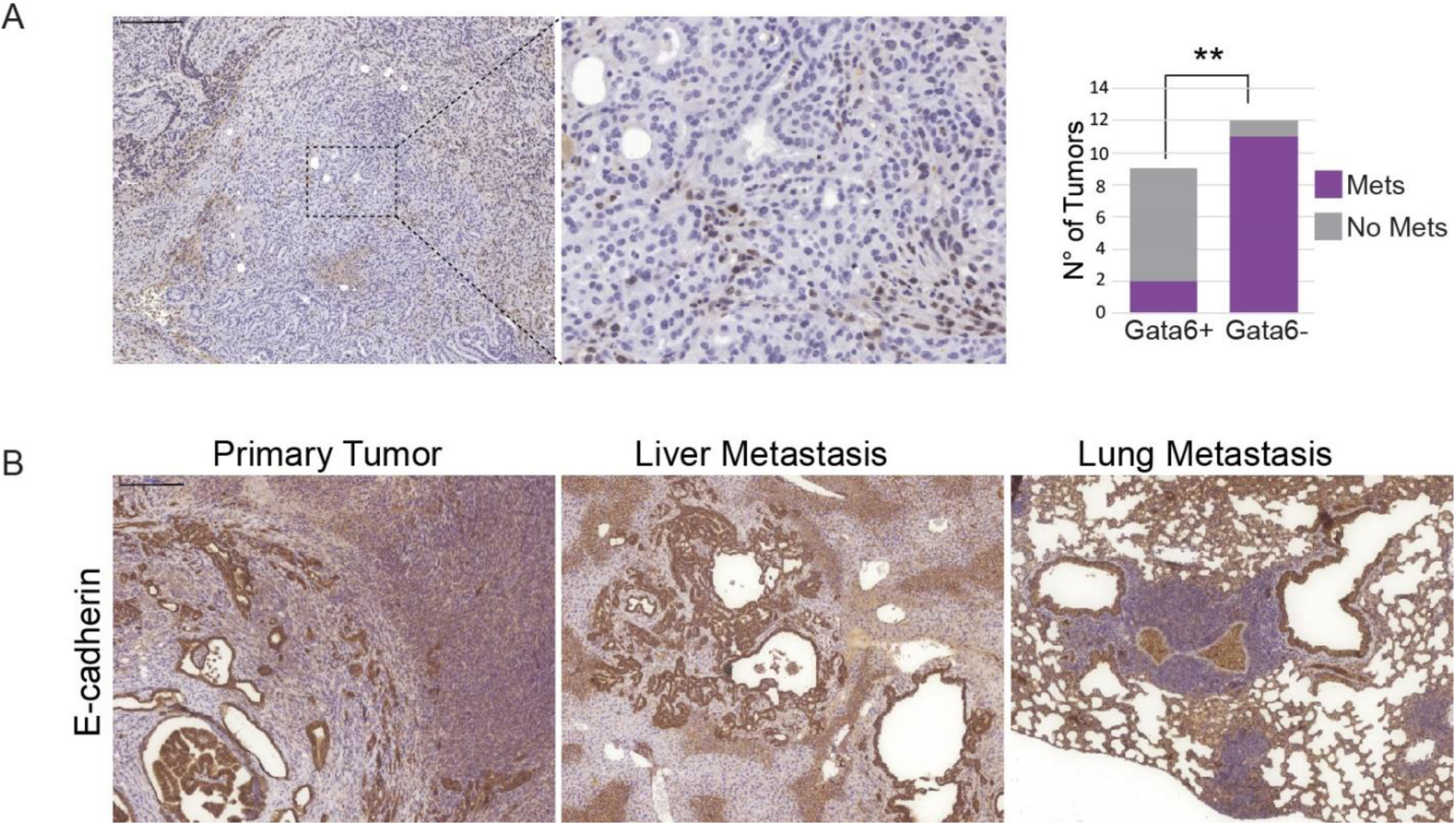
Spontaneous Gata6 loss favors metastasis. A) Representative images and quantification of Gata6 expression in primary tumors from an independent cohort of KFC mice. B) Representative images of E-cadherin expression in primary Gata6^LateKO^ tumors and the corresponding liver and lung metastases. Scale bar: 200µm.

**Supplementary Figure 6.**
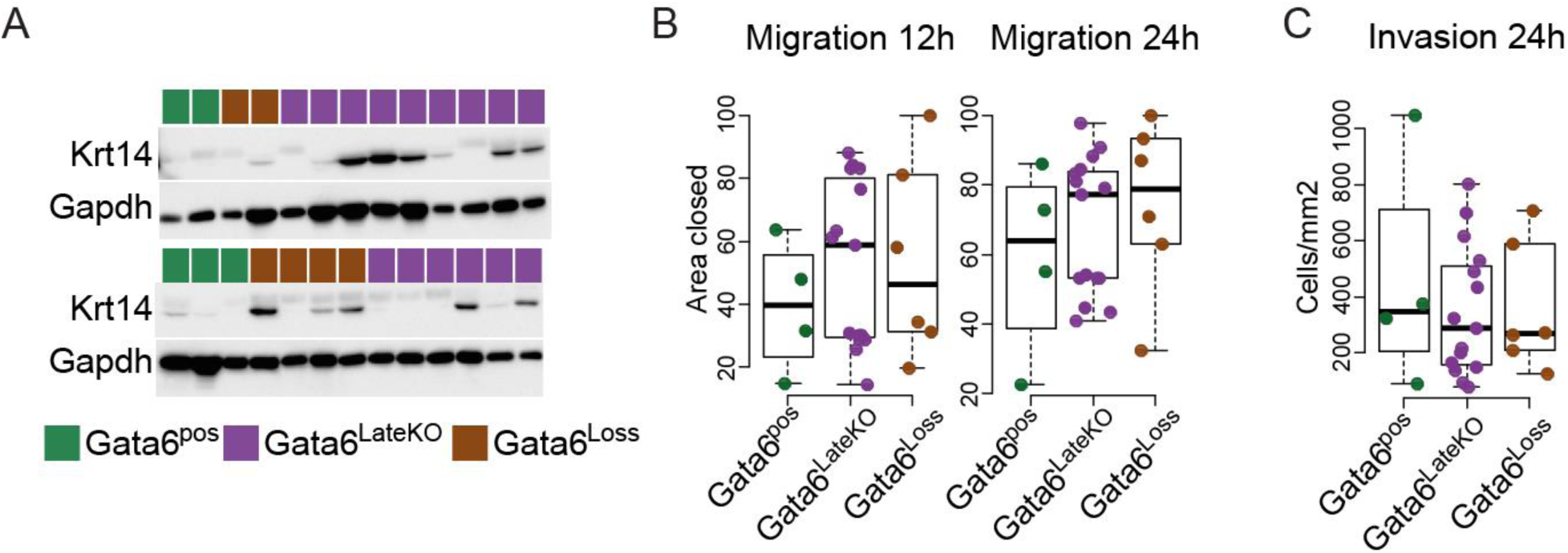
Characterization of primary KFC cells *in vitro*. A) Expression of Krt14 in primary KFC cells isolated from tumors of Gata6^pos^, Gata6^LateKO^ and Gata6^Loss^ mice, analyzed by western blotting from whole protein lysates. B) Quantification of *in vitro* migration of primary KFC cells, measured 12h and 24h after scratch. C) *In vitro* matrigel invasion assay with primary KFC cell lines, measured 24h after seeding. Gata6^pos^ n=4, Gata6^LateKO^ n=15, Gata6^Loss^ n=6.

**Supplementary Figure 7.**
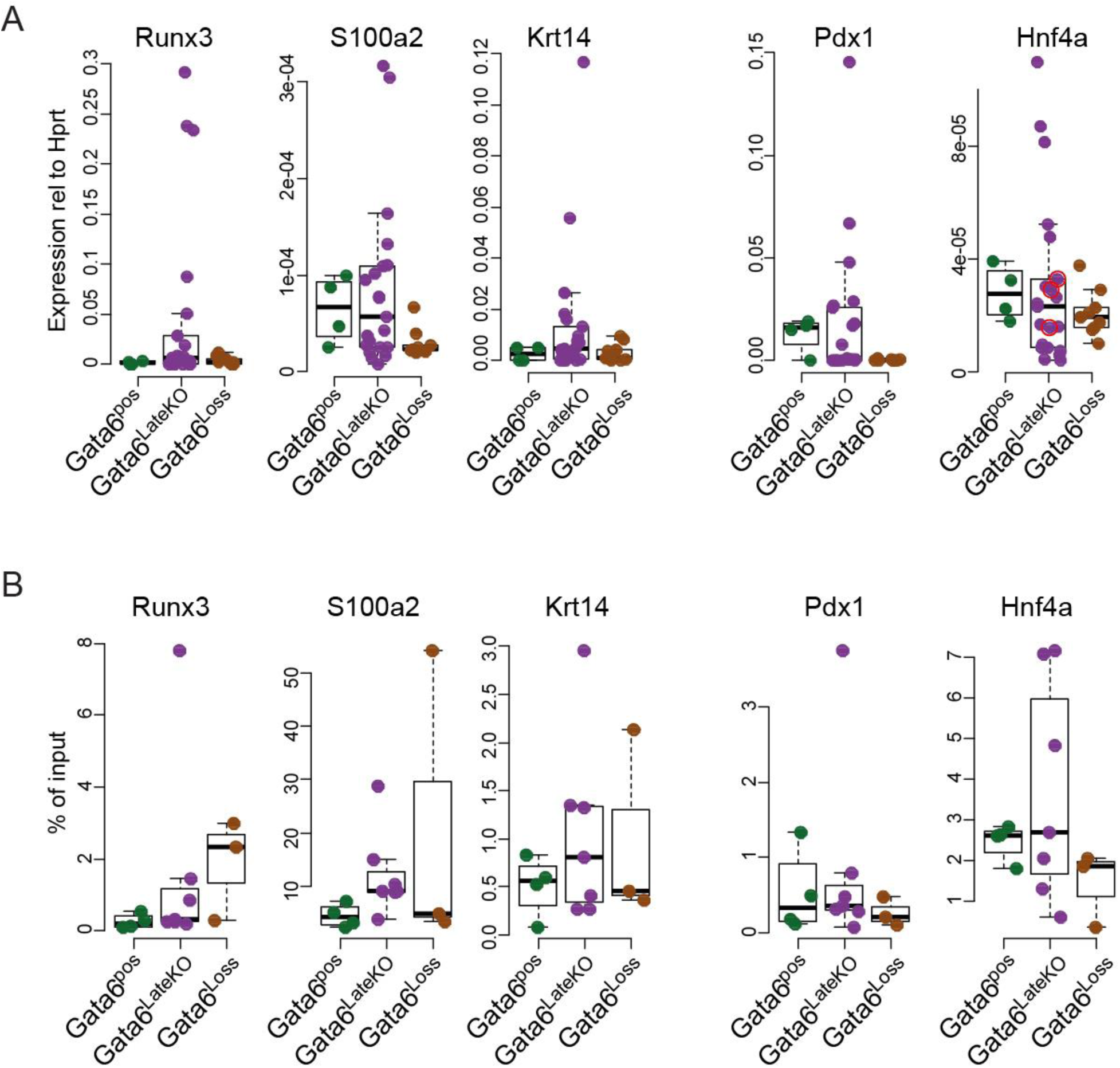
Gata6^neg^ KFC cells show features of basal PDAC. A) Expression of a set of basal and classical genes in Gata6^pos^, Gata6^LateKO^, and Gata6^Loss^ KFC cells measured by RT-qPCR. Gata6^pos^ n=4, Gata6^LateKO^ n=15, Gata6^Loss^ n=6. Each dot represents one cell line. B) H3K27ac binding to the promoter of Runx3, S100a2, Krt14, Pdx1 and Hnf4a, detected by ChIP-qPCR in primary KFC cells. Data are represented as % of input chromatin. Gata6^pos^ n=4, Gata6^LateKO^ n=7, Gata6^Loss^ n=3. Each dot represents the average value of at least three independent experiments for each tumor cell line.

**Supplementary Figure 8.**
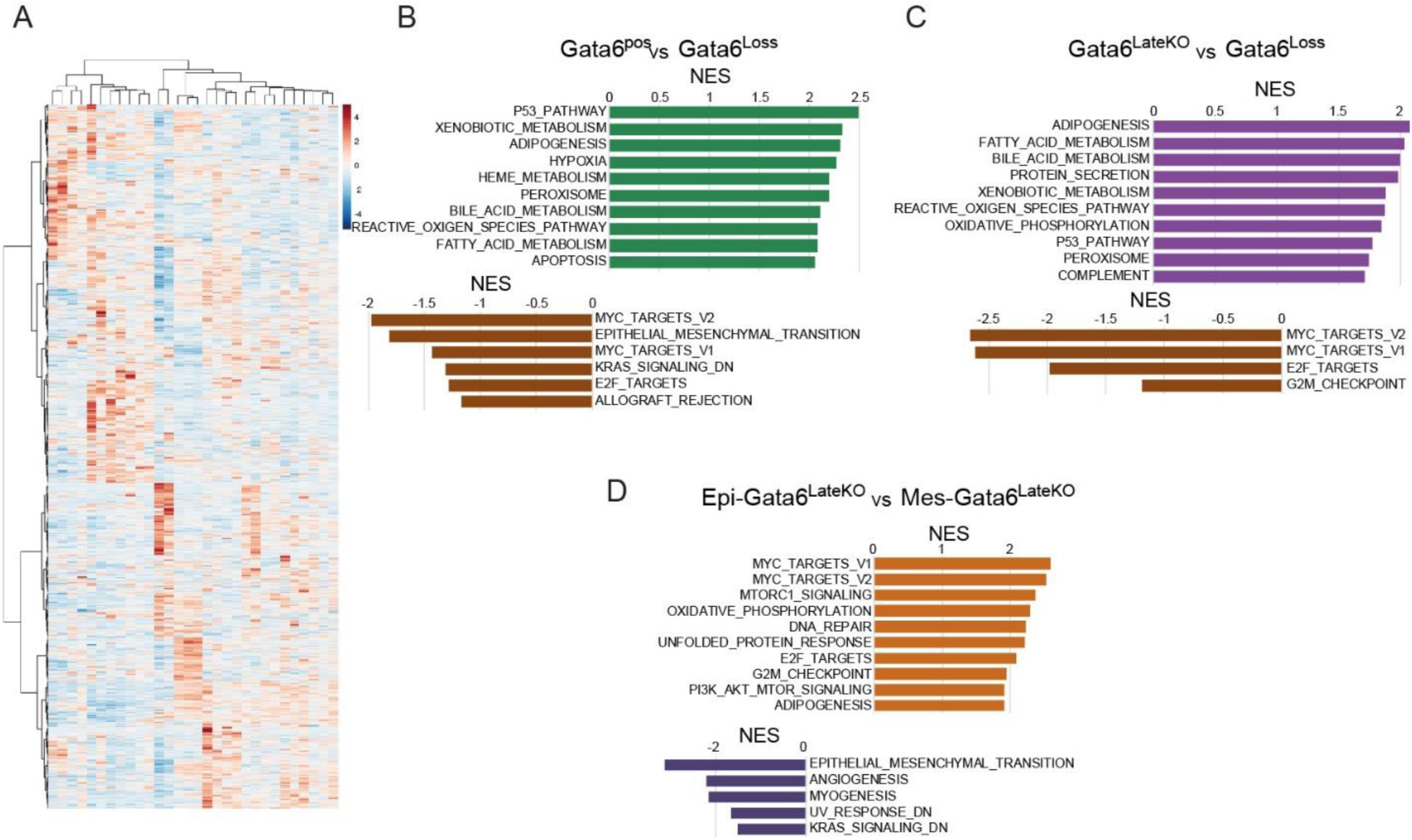
Gata6 loss induces cell plasticity. A) Heat map of unsupervised hierarchical clustering of Gata6^pos^, Gata6^LateKO^ and Gata6^Loss^ primary cells and reference cell lines. B-D) Gene set enrichment analyses of RNAseq data from KFC cells from the indicated subgroups.

**Supplementary Figure 9.**
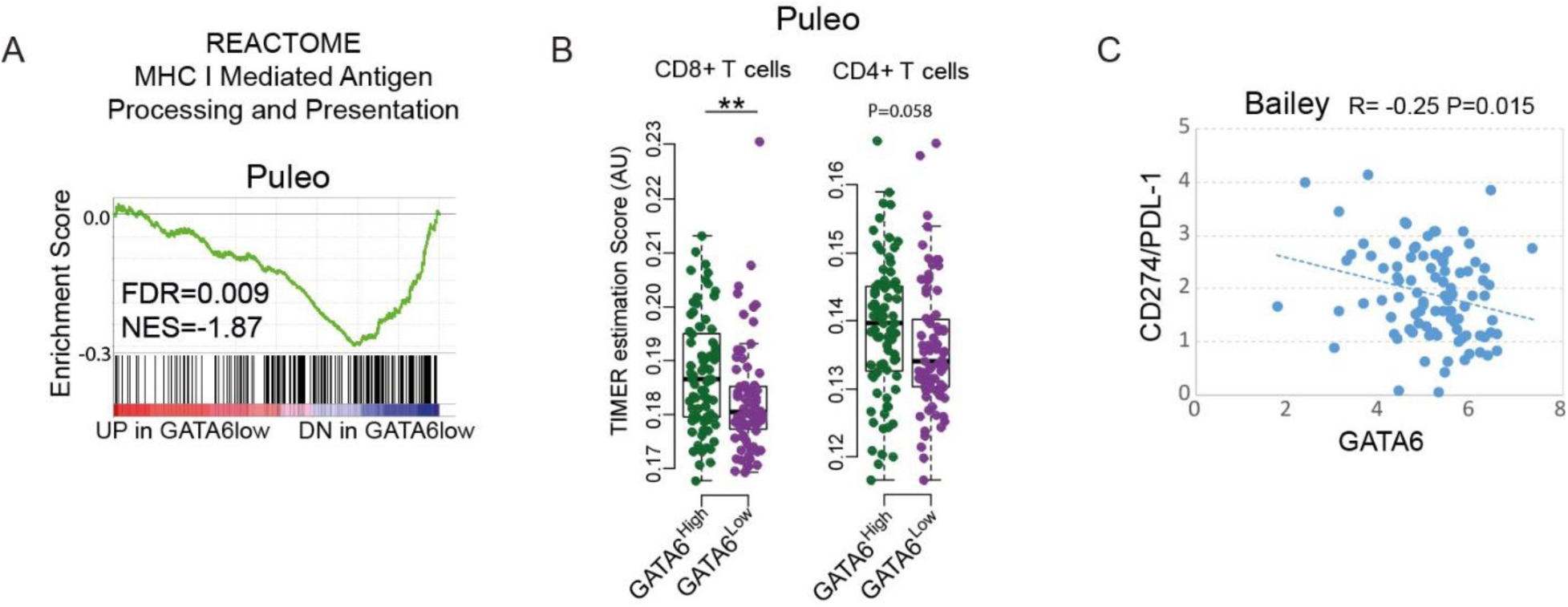
GATA6^low^ tumors show features of immune suppression. A) Estimated abundance of CD8+ and CD4+ T cells in GATA6^high^ and GATA6^low^ tumors of the Puleo dataset. **P<0.001. B) Correlation between GATA6 and CD274/PDL-1 expression in the Bailey dataset.

## References

1. Bailey P, Chang DK, Nones K, et al. Genomic analyses identify molecular subtypes of pancreatic cancer. Nature 2016;531(7592):47–52. doi: 10.1038/nature16965

2. Collisson EA, Sadanandam A, Olson P, et al. Subtypes of pancreatic ductal adenocarcinoma and their differing responses to therapy. Nat Med 2011;17(4):500–3. doi: 10.1038/nm.2344

3. Moffitt RA, Marayati R, Flate EL, et al. Virtual microdissection identifies distinct tumor-and stroma-specific subtypes of pancreatic ductal adenocarcinoma. Nat Genet 2015;47(10):1168–78. doi: 10.1038/ng.3398

4. Puleo F, Nicolle R, Blum Y, et al. Stratification of Pancreatic Ductal Adenocarcinomas Based on Tumor and Microenvironment Features. Gastroenterology 2018 doi: 10.1053/j.gastro.2018.08.033

5. Chan-Seng-Yue M, Kim JC, Wilson GW, et al. Transcription phenotypes of pancreatic cancer are driven by genomic events during tumor evolution. Nat Genet 2020 doi: 10.1038/s41588-019-0566-9 [published Online First: 2020/01/15]

6. Cancer Genome Atlas N. Comprehensive molecular portraits of human breast tumours. Nature 2012;490(7418):61–70. doi: 10.1038/nature11412

7. Martinelli P, Canamero M, del Pozo N, et al. Gata6 is required for complete acinar differentiation and maintenance of the exocrine pancreas in adult mice. Gut 2013;62(10):1481–8. doi: 10.1136/gutjnl-2012-303328

8. Martinelli P, Madriles F, Canamero M, et al. The acinar regulator Gata6 suppresses KrasG12V-driven pancreatic tumorigenesis in mice. Gut 2016;65(3):476–86. doi: 10.1136/gutjnl-2014-308042

9. Martinelli P, Carrillo-de Santa Pau E, Cox T, et al. GATA6 regulates EMT and tumour dissemination, and is a marker of response to adjuvant chemotherapy in pancreatic cancer. Gut 2017;66(9):1665–76. doi: 10.1136/gutjnl-2015-311256

10. Aung KL, Fischer SE, Denroche RE, et al. Genomics-Driven Precision Medicine for Advanced Pancreatic Cancer - Early Results from the COMPASS Trial. Clin Cancer Res 2017 doi: 10.1158/1078-0432.CCR-17-2994

11. Hamdan FH, Johnsen SA. DeltaNp63-dependent super enhancers define molecular identity in pancreatic cancer by an interconnected transcription factor network. Proc Natl Acad Sci U S A 2018;115(52):E12343–E52. doi: 10.1073/pnas.1812915116 [published Online First: 2018/12/14]

12. Somerville TDD, Xu Y, Miyabayashi K, et al. TP63-Mediated Enhancer Reprogramming Drives the Squamous Subtype of Pancreatic Ductal Adenocarcinoma. Cell Rep 2018;25(7):1741–55 e7. doi: 10.1016/j.celrep.2018.10.051 [published Online First: 2018/11/15]

13. Andricovich J, Perkail S, Kai Y, et al. Loss of KDM6A Activates Super-Enhancers to Induce Gender-Specific Squamous-like Pancreatic Cancer and Confers Sensitivity to BET Inhibitors. Cancer Cell 2018;33(3):512–26 e8. doi: 10.1016/j.ccell.2018.02.003

14. Lomberk G, Blum Y, Nicolle R, et al. Distinct epigenetic landscapes underlie the pathobiology of pancreatic cancer subtypes. Nat Commun 2018;9(1):1978. doi: 10.1038/s41467-018-04383-6 [published Online First: 2018/05/19]

15. Schaefer CF, Anthony K, Krupa S, et al. PID: the Pathway Interaction Database. Nucleic Acids Res 2009;37(Database issue):D674–9. doi: 10.1093/nar/gkn653 [published Online First: 2008/10/04]

16. Sodhi CP, Li J, Duncan SA. Generation of mice harbouring a conditional loss-of-function allele of Gata6. BMC Dev Biol 2006;6:19. doi: 10.1186/1471-213X-6-19 [published Online First: 2006/04/14]

17. Schonhuber N, Seidler B, Schuck K, et al. A next-generation dual-recombinase system for time- and host-specific targeting of pancreatic cancer. Nat Med 2014;20(11):1340–47. doi: 10.1038/nm.3646

18. Chen Y, LeBleu VS, Carstens JL, et al. Dual reporter genetic mouse models of pancreatic cancer identify an epithelial-to-mesenchymal transition-independent metastasis program. EMBO Mol Med 2018;10(10) doi: 10.15252/emmm.201809085 [published Online First: 2018/08/19]

19. Iacobuzio-Donahue CA, Fu B, Yachida S, et al. DPC4 gene status of the primary carcinoma correlates with patterns of failure in patients with pancreatic cancer. J Clin Oncol 2009;27(11):1806–13. doi: 10.1200/JCO.2008.17.7188 [published Online First: 2009/03/11]

20. Ocana OH, Corcoles R, Fabra A, et al. Metastatic colonization requires the repression of the epithelial-mesenchymal transition inducer Prrx1. Cancer Cell 2012;22(6):709–24. doi: 10.1016/j.ccr.2012.10.012 [published Online First: 2012/12/04]

21. Tsai JH, Donaher JL, Murphy DA, et al. Spatiotemporal regulation of epithelial-mesenchymal transition is essential for squamous cell carcinoma metastasis. Cancer Cell 2012;22(6):725–36. doi: 10.1016/j.ccr.2012.09.022 [published Online First: 2012/12/04]

22. Reichert M, Bakir B, Moreira L, et al. Regulation of Epithelial Plasticity Determines Metastatic Organotropism in Pancreatic Cancer. Dev Cell 2018;45(6):696–711 e8. doi: 10.1016/j.devcel.2018.05.025 [published Online First: 2018/06/20]

23. McDonald OG, Li X, Saunders T, et al. Epigenomic reprogramming during pancreatic cancer progression links anabolic glucose metabolism to distant metastasis. Nat Genet 2017;49(3):367–76. doi: 10.1038/ng.3753 [published Online First: 2017/01/17]

24. Mueller S, Engleitner T, Maresch R, et al. Evolutionary routes and KRAS dosage define pancreatic cancer phenotypes. Nature 2018;554(7690):62–68. doi: 10.1038/nature25459

25. Li T, Fan J, Wang B, et al. TIMER: A Web Server for Comprehensive Analysis of Tumor-Infiltrating Immune Cells. Cancer Res 2017;77(21):e108–e10. doi: 10.1158/0008-5472.CAN-17-0307 [published Online First: 2017/11/03]

26. Muckenhuber A, Berger AK, Schlitter AM, et al. Pancreatic Ductal Adenocarcinoma Subtyping Using the Biomarkers Hepatocyte Nuclear Factor-1A and Cytokeratin-81 Correlates with Outcome and Treatment Response. Clin Cancer Res 2018;24(2):351–59. doi: 10.1158/1078-0432.CCR-17-2180 [published Online First: 2017/11/05]

27. Noll EM, Eisen C, Stenzinger A, et al. CYP3A5 mediates basal and acquired therapy resistance in different subtypes of pancreatic ductal adenocarcinoma. Nat Med 2016;22(3):278–87. doi: 10.1038/nm.4038 [published Online First: 2016/02/09]

28. Kalisz M, Bernardo E, Beucher A, et al. HNF1A recruits KDM6A to activate differentiated acinar cell programs that suppress pancreatic cancer. EMBO J 2020:e102808. doi: 10.15252/embj.2019102808 [published Online First: 2020/03/11]

29. Molero X, Vaquero EC, Flandez M, et al. Gene expression dynamics after murine pancreatitis unveils novel roles for Hnf1alpha in acinar cell homeostasis. Gut 2012;61(8):1187–96. doi: 10.1136/gutjnl-2011-300360 [published Online First: 2011/09/29]

30. Aiello NM, Maddipati R, Norgard RJ, et al. EMT Subtype Influences Epithelial Plasticity and Mode of Cell Migration. Dev Cell 2018;45(6):681–95 e4. doi: 10.1016/j.devcel.2018.05.027 [published Online First: 2018/06/20]

31. Manguso RT, Pope HW, Zimmer MD, et al. In vivo CRISPR screening identifies Ptpn2 as a cancer immunotherapy target. Nature 2017;547(7664):413–18. doi: 10.1038/nature23270 [published Online First: 2017/07/21]

32. Burr ML, Sparbier CE, Chan KL, et al. An Evolutionarily Conserved Function of Polycomb Silences the MHC Class I Antigen Presentation Pathway and Enables Immune Evasion in Cancer. Cancer Cell 2019;36(4):385–401 e8. doi: 10.1016/j.ccell.2019.08.008 [published Online First: 2019/10/01]

33. Cobo I, Martinelli P, Flandez M, et al. Transcriptional regulation by NR5A2 links differentiation and inflammation in the pancreas. Nature 2018;554(7693):533–37. doi: 10.1038/nature25751 [published Online First: 2018/02/15]

## References

3. Moffitt RA, Marayati R, Flate EL, et al. Virtual microdissection identifies distinct tumor- and stroma-specific subtypes of pancreatic ductal adenocarcinoma. Nat Genet 2015;47(10):1168–78. doi: 10.1038/ng.3398

25. Li T, Fan J, Wang B, et al. TIMER: A Web Server for Comprehensive Analysis of Tumor-Infiltrating Immune Cells. Cancer Res 2017;77(21):e108-e10. doi: 10.1158/0008-5472.CAN-17-0307 [published Online First: 2017/11/03]

34. Collier LS, Carlson CM, Ravimohan S, et al. Cancer gene discovery in solid tumours using transposon-based somatic mutagenesis in the mouse. Nature 2005;436(7048):272–6. doi: 10.1038/nature03681 [published Online First: 2005/07/15]

35. Perez-Mancera PA, Rust AG, van der Weyden L, et al. The deubiquitinase USP9X suppresses pancreatic ductal adenocarcinoma. Nature 2012;486(7402):266–70. doi: 10.1038/nature11114 [published Online First: 2012/06/16]

36. Parekh S, Ziegenhain C, Vieth B, et al. The impact of amplification on differential expression analyses by RNA-seq. Sci Rep 2016;6:25533. doi: 10.1038/srep25533 [published Online First: 2016/05/10]

37. Macosko EZ, Basu A, Satija R, et al. Highly Parallel Genome-wide Expression Profiling of Individual Cells Using Nanoliter Droplets. Cell 2015;161(5):1202–14. doi: 10.1016/j.cell.2015.05.002 [published Online First: 2015/05/23]

